# The orbitofrontal cortex forms a context-generalized spatial schema that preserves topology and distance

**DOI:** 10.1101/2025.09.15.676202

**Authors:** Raunak Basu, Marjan Mozaffarilegha, Ipek Bölükbaşı, Cansu Üstüner, Hiroshi T. Ito

## Abstract

Flexible and efficient navigation requires the brain to construct maps that are both topological, preserving the relationships between locations, and schematic, enabling generalization across environments. Although spatial maps in the hippocampus (HPC) and medial entorhinal cortex (MEC) have been extensively studied, they remap almost orthogonally across environments even during identical behaviors, raising the question of how animals maintain consistent navigation strategies across spatial contexts. Here, we identify a novel spatial map in the orbitofrontal cortex (OFC) that encodes navigational targets with distinct neural representations while preserving their topological order and relative distances by scaling to physical path lengths. Remarkably, OFC maps remained stable when animals performed the same navigation task across rooms and maze geometries, in stark contrast to the pronounced remapping observed in HPC and MEC. Moreover, OFC maps persisted after HPC or MEC lesions, demonstrating an independent spatial mapping system. These findings reveal a task-relevant topological schema in the OFC that uniquely supports flexible context-invariant navigation, expanding the brain’s spatial mapping repertoire beyond the hippocampal–entorhinal system.

## INTRODUCTION

Animals, including rodents and humans, navigate flexibly within and across environments by leveraging the knowledge of spatial structures abstracted from landmark relationships and previously learned behavioral strategies^1^. For instance, when revisiting your hometown after many years, several landmarks may have changed, yet core spatial relationships, such as the location of your home relative to the train station, typically remain stable. This preserved relational structure enables efficient navigation with only partial updates to existing knowledge, eliminating the need to re-learn the entire environment from scratch. Such reuse of pre-existing spatial memory structures^2^ allows the brain to interpret novel experiences through the lens of prior knowledge, which is a critical capacity in real-world settings where identical situations rarely repeat.

Extensive research has established the hippocampus (HPC) and medial entorhinal cortex (MEC) as crucial nodes in the neural circuits supporting spatial memory and navigation. In the HPC, place cells fire in a location-specific manner^3^, while in the MEC, grid cells exhibit periodic firing fields that tile the entire environment in a hexagonal pattern^4^. As an animal traverses through space, ensemble neural activity in HPC and MEC changes continuously and smoothly. As a result, at the neural population level, both place cells and grid cells form a topology-preserved map of an environment where nearby locations are mapped through similar ensemble firing patterns^5–8^. However, these spatial representations are highly context-specific. Manipulations such as changing the environmental shape^9^, room^9,10^, or surrounding cues^11^, or even altering the task rules^12,13^ induce ‘global remapping’ in HPC and MEC neurons, where firing fields shift or disappear in one of the contexts. This remapping leads to the formation of orthogonal spatial maps between contexts. For example, grid cell firing patterns shift in response to the presence of novel landmarks, resulting in distinct population-level representations even for the same physical space^11^. Hence, while HPC and MEC neurons encode detailed spatial context and topology, their representations are not likely suited for generalizing spatial knowledge across different environmental contexts.

By contrast, behavioral studies have demonstrated that rodents can generalize learned rules and behaviors to new contexts, and recent evidence implicates the prefrontal cortex as a key structure for this ability. For example, in a task requiring animals to visit spatial positions in a fixed sequence, rodents quickly adapt to new sequences composed of different positions. Here, prefrontal neurons encode the ordinal structure of the sequence by generalizing across location identities^14^. Similarly, in an olfactory sequence task, animals rapidly adapt to new sets of odor cues composing the sequence, which is reflected in the neural activity in OFC – subregion of the prefrontal cortex – representing the sequence structure by generalizing across odor identities^15,16^. Generalization of spatial learning and navigation strategies has also been well documented. Rodents trained to navigate to a goal in one environment can rapidly do so even when the entire setup is relocated to a different room^17^. Another study demonstrated that rats trained on a set of cue-place associations are able to learn novel cue-place pairings more quickly than during initial acquisition^2^. As a candidate neural mechanism supporting this assimilation of novel cue–place associations into pre-existing spatial memory frameworks, studies involving brain lesions and immediate early gene expressions have identified the prefrontal cortex as a crucial region^18,19^. These previous studies collectively highlight the prefrontal cortex as a likely key locus for spatial memory schemas that enable knowledge transfer across environments. Yet, the neural code underlying this capacity – specifically how prefrontal neural ensembles abstract spatial structure across environments while preserving topological relationship between locations, and how this process differs from the environment-specific maps in HPC and MEC – remains elusive, leaving a critical gap in our understanding of the neural basis of context-generalized spatial schemas.

Here, we address this question by characterizing the neural coding scheme in OFC underlying the generalization of spatial knowledge across environments. In our previous work, we developed a novel OFC-dependent goal-directed navigation task in an environment with multiple reward locations, where OFC neural ensembles represent the animal’s future spatial goal even before the onset of the journey^20^. This task paradigm is particularly well-suited to disentangle spatial coding in the OFC from its plausible representation of task structure. Prefrontal neurons are known to represent progress towards the goal^14^, which can be confounded with spatial position unless the animal navigates toward different goals in the same maze^21,22^. By introducing frequent goal switches, our task design overcomes this limitation and enables the identification of true spatial representations in the OFC. Applying this task paradigm across 3 mazes with different geometric shapes and lengths, we now show that the OFC spatial representations distinguish individual reward locations while preserving both topological and distance relationships across environments. To our knowledge, this represents the first example of a topology-preserved spatial map outside the hippocampal and parahippocampal regions. But importantly, the OFC map differs in key ways from previously described spatial maps. To assess the generalization ability of these maps across contexts, we trained animals to perform the same task in either a novel room or a maze with a different geometric shape while recording from OFC, HPC, and MEC. In both cases, while HPC and MEC ensembles underwent global remapping, OFC activity preserved its spatial structure, indicating robust context generalization. Furthermore, by lesioning HPC and MEC, we found that the formation of OFC spatial map occurs independently of HPC and MEC maps. Our results identify a previously unrecognized spatial mapping system in the OFC that encodes task-relevant topological relationships and supports schema-based generalization of spatial knowledge across environments, playing a distinct and complementary role in spatial memory and navigation alongside the hippocampal–entorhinal system.

## RESULTS

### The OFC exhibits spatial coding dedicated to navigational targets

To compare the spatial coding properties of OFC neural ensembles with those in HPC and MEC, we leveraged the goal-directed navigation task comprising ten equally spaced reward wells, developed in our previous work^20^. The task required animals to alternate between designated pairs of wells, receiving a water reward upon identifying the correct well (Figure 1A). After five or more consecutive correct trials, the goal well pair was switched, which required the animals to update their navigational goals accordingly. Once trained, we implanted tetrode microdrives targeting the OFC (n = 9 rats; mean: 157 neurons/session; range: 68-328 neurons), hippocampal CA1 region (n = 3 rats; mean: 73 neurons/session; range: 53-98 neurons), or the MEC (n = 2 rats; mean: 67 neurons/session; range: 53-83 neurons; Figure S1).

**Figure 1:**
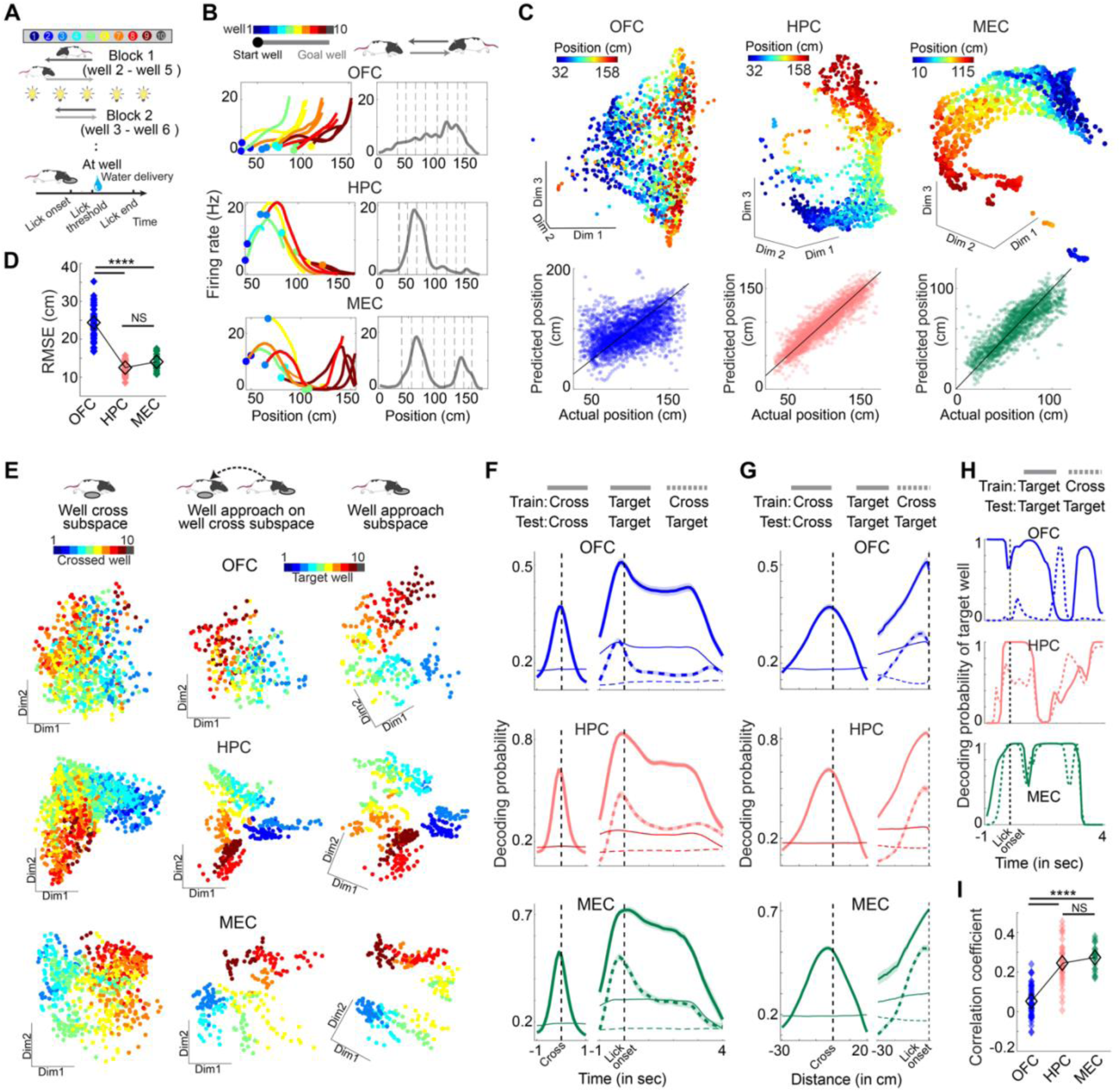
OFC encodes discrete navigational targets in contrast to continuous spatial mapping in HPC and MEC. **(A)** Schematic of the task (top) and the sequence of events during licking a reward well (bottom). **(B)** Firing rates of a representative neuron from OFC (top), HPC (middle), and MEC (bottom) as a function of position. Left panel shows trial-averaged firing rates for each start-goal well combination, represented with a distinct color-code (circle color represents the start location, and line color represents the goal location). Right panel shows the mean firing rates across the maze position. Dotted vertical lines represent well locations (wells 2-9 were used). Firing rates for only one movement direction are plotted. **(C)** (top) UMAP projections of neural ensemble activity from OFC (left), HPC (middle), and MEC (right) for a representative session and color-coded based on the animal’s location. Only journeys in one movement direction are shown. Bottom panel plots the animal’s decoded position from ensemble neural activity during periods of motion against its actual location within the corresponding session. The identity lines are shown in black. Each dot represents data from a 100-ms time bin. **(D)** Root mean square errors (RMSEs) of the animal’s instantaneous position decoded from neural ensembles in the OFC, HPC, and MEC. **(E)** Neural ensemble activity in a representative session each, from OFC (top), HPC (middle), and MEC (bottom), during crossings of non-targeting wells (left) and approaches to target wells (middle and left), projected on the first two orthogonalized LDA dimensions. Middle panels show neural activity during approaches to target wells, projected on LDA dimensions trained on non-targeting wells during crossing (i.e., the subspace computed in the left panel), while right panels depict those projected on LDA dimensions trained on targeting wells. For each trial, three time points (300 milliseconds) leading up to crossing (for left panels) and licking (middle and right panels) are plotted. Note the similarity in the target-well approach representations in the middle and right panels for HPC and MEC, but not OFC. **(F)** Decoding probabilities for non-targeting wells during crossings (solid line on left,-1 to 1 s relative to crossing) and for target wells during approach and subsequent licking. Decoders were trained either on non-target wells (Cross: thick dotted line on right) or target wells (Target: solid line on right,-1 to 4 sec relative to lick onset). Analyses were performed on neural ensembles from OFC (top), HPC (middle), and MEC (bottom). **(G)** Same as **F**, but decoding results are plotted against distance relative to the decoding wells (Cross well: 30 cm prior to crossing to 20 cm post crossing; Target well: 30 cm prior to arrival). In panels **F** and **G**, means (solid and thick dotted) ± s.e.m. (shading) are shown. Thin colored lines indicate chance levels. **(H)** Decoding probability of target wells using either the cross-well decoder (dotted) or the target-well decoder (solid), shown for-1 to 4 s relative to lick onset in a representative trial. **(I)** Correlation coefficients between decoding probabilities obtained from the cross-well and target-well decoders. Each diamond represents the average correlation coefficient across all trials within a session. In panels **D** and **I**, filled colored and open black diamonds represent individual sessions and their mean, respectively. ****p < 0.0001

Because spatial representations have been typically studied during locomotion, we first focused our analysis on spatial coding as animals actively navigated toward various target wells. As expected from previous studies, HPC and MEC neurons exhibited increased firing when the animal visited specific locations in the linear maze. In contrast, OFC neurons showed broader and less position-specific firing over the maze (Figures 1B and S2). To further understand how different spatial positions along the linear maze are represented in simultaneously recorded neural ensembles, we applied Uniform Manifold Approximation and Projection (UMAP)^23^ on individual 100-ms bins of neural population activity during navigation. This analysis revealed that HPC and MEC neural manifolds encode the maze positions as a continuum, with nearby locations represented by similar ensemble firing patterns. In contrast, such a spatial structure of neural activity between locations was largely missing in the OFC (Figure 1C top). To quantify these differences in spatial coding, we applied Gaussian process regression (GPR) with a squared exponential kernel to decode the animal’s instantaneous location from neural population activity. The GPR enables regression of continuous variables (like position) that smoothly vary over a manifold, and can accommodate nonlinear relationships, such as those reported in HPC and MEC population manifolds^6–8^. Consistent with the UMAP results and the spatial tuning properties of individual neurons, the GPR-based decoding of the animal’s location was significantly better for HPC and MEC than for OFC (Figures 1C bottom and 1D; root mean squared error, mean ± s.e.m.: 24.38 ± 0.5 cm from n = 62 sessions for OFC, 12.5 ± 0.26 cm from n = 37 sessions for HPC, and 14.03 ± 0.44 cm from n = 21 session for MEC). Overall, neural ensembles in HPC and MEC, but not OFC, robustly encode the animal’s location during locomotion.

However, in our previous work, we found that OFC neurons encode the identities of wells specifically when an animal approaches and licks them^20^, suggesting that spatial coding in the OFC is primarily dedicated to behaviorally-relevant navigational targets in the maze. This unique goal-oriented coding scheme led us to ask if HPC and MEC similarly form dedicated maps for navigational targets or whether their spatial codes remain invariant across two distinct navigational states - simply running over or targeting a specific location. To address this question, we compared neural representations between the two states, either when the animal traversed one of the wells (“cross-well activity”) or when it approached the well as a navigational target (“target-well activity”). For the cross-well events, we projected 300 ms of neural activity preceding each crossing into a subspace optimized to separate the cross-well identities (“cross-well subspace” using linear discriminant analysis, LDA; see Methods). Similarly, for target-well events, we analyzed 300 ms of neural activity preceding lick onset at a target well, projecting it either on the cross-well subspace or on another subspace optimized for separating target-well identities (“target-well subspace”, Figure 1E). If the same spatial mapping scheme is shared between cross-and target-wells, the neural representations of well identities should look similar between the two corresponding subspaces. This was indeed the case in both HPC and MEC, where well-specific activities for target-wells resembled those for cross-wells. In contrast, OFC neurons exhibited well-specific coding only when projecting on the target-well subspace (Figure 1E).

To quantify these observations, we trained two types of well-identity decoders, one for non-licked wells traversed during navigation (cross-well decoder; trained on-0.2 to 0s relative to crossing) and the other for target wells where the animal approached and licked (target-well decoder; trained on-0.5s to 3s relative to lick onset). To account for the distinct spatial maps for the two different running directions^24^, we used a Gaussian mixture model based decoder^25^ that afforded two subclasses to each well (Methods). We then tested whether the cross-well decoder could generalize to predict the identity of target wells. This was expected when spatial coding remains consistent across navigational states, and we indeed found that the cross-well decoder performance on target wells was significantly higher in HPC and MEC than in OFC (Figures 1F-G; mean ± s.e.m. of peak target-well decoding probabilities: 0.287 ± 0.008 in OFC; 0.491 ± 0.019 in HPC; 0.507± 0.017 in MEC; p = 1.13×10^-13^ OFC versus HPC, p = 2.89×10^-11^ OFC versus MEC, p = 0.85 HPC versus MEC by Kruskal-Wallis test followed by pairwise comparison). The results remained the same even after controlling for differences in decoding strength across brain regions by using subsampled datasets (Figure S2). Moreover, we confirmed that the OFC’s selective spatial coding for navigational targets is not merely driven by reward, because well-specific representations in the OFC emerged prior to the onset of reward delivery (Figure S3), suggesting that the OFC’s goal-directed spatial map forms as the animal plans to approach a specific location. Together, these findings confirmed that, while HPC and MEC maintain consistent spatial coding across navigational states, OFC forms a spatial representation that selectively emphasizes behaviorally-relevant locations targeted by the animal.

Finally, to assess the similarity of spatial coding between cross-and target-wells at the single trial level, we analyzed temporal correlations between the cross-well and target-well decoder performances in predicting target-well identity. In both HPC and MEC, target-well decoding probabilities between the two decoders were significantly more correlated than in OFC (Figures 1H-I; mean ± s.e.m. of correlation coefficients: 0.052 ± 0.009 in OFC; 0.246 ± 0.017 in HPC; 0.274 ± 0.014 in MEC; p = 3.82×10^-12^ OFC versus HPC, p = 2.24×10^-11^ OFC versus MEC, p = 0.62 HPC versus MEC by Kruskal-Wallis test followed by pairwise comparison), further supporting the idea of consistent spatial coding across navigational states in HPC and MEC, but not in OFC. These findings persisted even when controlling for differences in decoding strength by using subsampled datasets or by using an alternative time window (1.1 seconds centered at lick onset) during which the decoding performances were typically highest (Figure S2).

Taken together, our results reveal a distinct spatial coding scheme in the OFC that selectively maps behaviorally-relevant locations targeted by the animal, but not other locations traversed during journeys. In contrast, HPC and MEC neurons maintain largely consistent spatial maps regardless of the navigational state.

### The OFC spatial map preserves the topological arrangement of encoded positions

A hallmark of spatial maps is the preservation of the topological and distance relationships between locations. Hence, nearby locations should be represented by similar neural activity, while neural dissimilarity increases proportionally with physical distance. To test if OFC neurons form a spatial map that preserves topological relationships, we focused on a window spanning 0.5 s prior (well-approaching) to 3 s after the onset of licking. This window was chosen as the decoding probability of target-well identity peaks just prior to or at the lick onset in all three brain regions (Fig 1F, median peak probability: 0.1 s, 0.1 s, and 0 s prior to lick onset for OFC, HPC, and MEC respectively), and remains high through the entire lick duration^20^ (median lick duration: 3.31 s, n = 13946 rewarded lick events in the linear maze). We found that, during well-approach and licking phases, individual OFC neurons changed their overall firing rates depending on well identity, apparently reflecting the topological arrangement of well positions along the linear maze (Figures 2A, S3). This pattern was less visible in neurons in HPC and MEC, likely due to their sharp spatial tuning to specific locations in the maze (Figures 2A, S3). However, individual neurons often did not maintain activity across all well locations in the maze, nor did their firing rates vary monotonously along well positions, thereby precluding clear evidence of topological mapping along the linear maze at the single neuron level.

**Figure 2:**
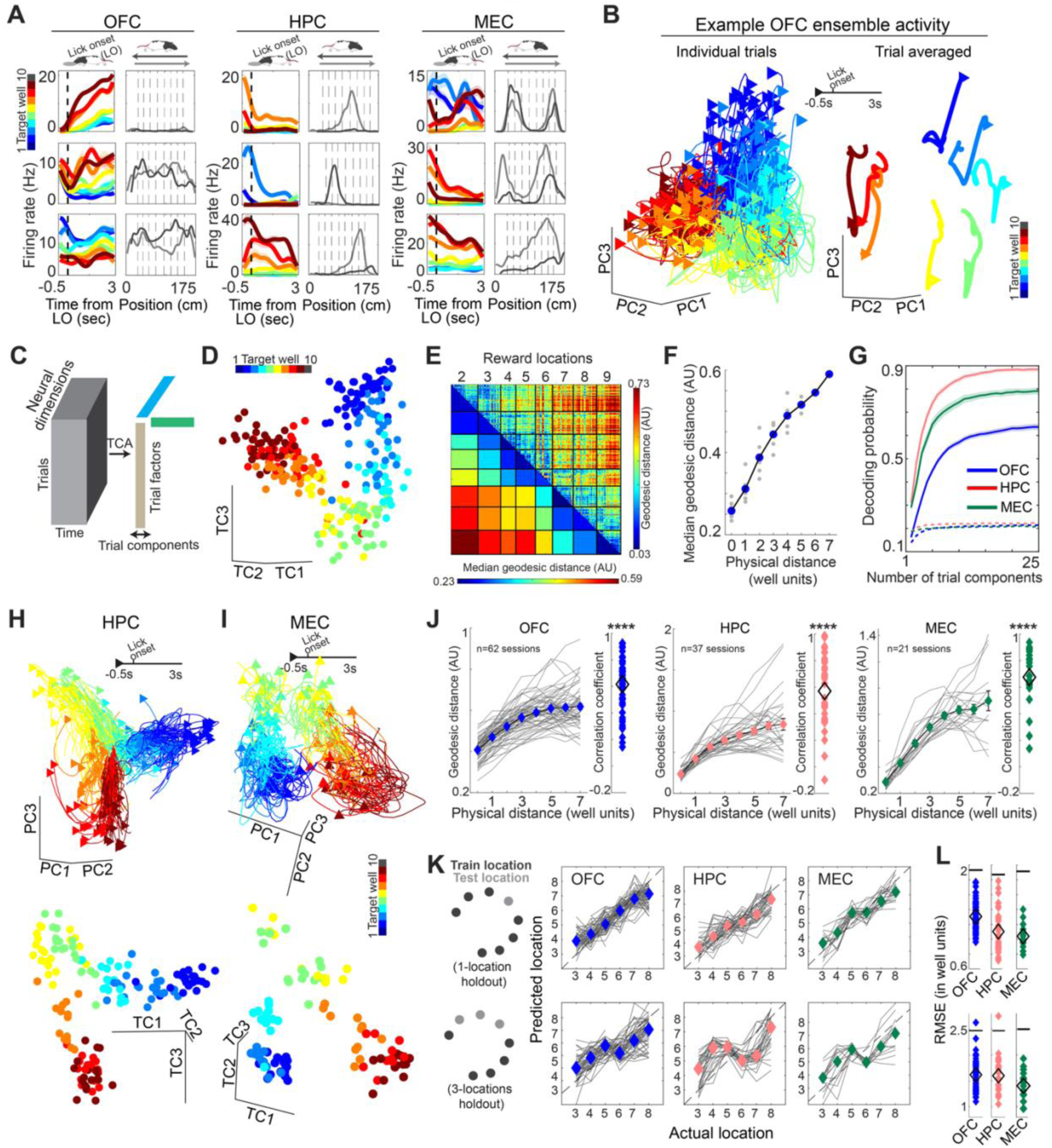
Topology preserved spatial maps in OFC, HPC, and MEC. **(A)** Firing rates of three representative neurons from OFC (left), HPC (middle), and MEC (right). For each neuron, the left panel shows firing rates averaged for individual target wells from 500 ms before to 3 s after lick onset. Right panel shows the mean firing rates along the maze positions with the two running directions plotted separately (light and dark grey). Dotted vertical lines represent well locations (wells 2-9 were used). **(B)** (left) OFC neural ensemble activity from an example session projected on the first three principal components. Each thin line depicts a single trial from 500 ms before to 3 s after the lick onset with color-code based on the well identity. Right panel shows the averaged activity for individual target wells. **(C)** Schematic of tensor component analysis (TCA). **(D)** Trial factors from the first three trial components were computed in the same session as in **B**. Each dot represents a single trial factor colored by the target well identity. **(E)** Geodesic distance between 8-dimensional trial factors from the same example session as **B**. Each pixel in the upper triangular part corresponds to the geodesic distance between two trial factors computed for a pair of target wells on an individual trial basis. Trial factors are grouped based on target well identity. The lower triangular part shows the median distance between each pair of target wells. **(F)** Median geodesic distances plotted against the physical distances between wells (in well units). Each gray circle represents a median distance (as in the lower triangular part of panel **E**) for a corresponding well distance pair, with their mean denoted by a blue circle. **(G)** Decoding probabilities of target wells against the number of trial components. Note that spatial information is primarily contained in the first few trial components in all three brain regions. Plotted are means (solid and thick dotted) ± s.e.m. (shading). Dotted lines show chance levels. **(H)** (top) HPC ensemble activity from an example session projected on the first three principal components (top) or on the first three TCA trial components (bottom). **(I)** Same as in **H**, but for MEC ensemble activity from an example session. Plot and color schemes in **H** and **I** are identical to those in panels **B** left and **D. (J)** Geodesic distances from OFC (left), HPC (middle), and MEC (right) ensembles plotted against physical distances in well units. Each gray curve represents the mean geodesic distance in a single session (as the black curve in panel **F**) with the means of all sessions denoted by diamonds. In each panel, the inset on the right shows the correlation coefficient between median geodesic distance and physical distance for individual sessions (colored diamonds) with their means across sessions denoted by diamonds. ****p < 0.0001 by Wilcoxon signed-rank test. (**K)** Prediction of target-well identities by Gaussian process regression (GPR) from the trial factors obtained from OFC (left), HPC (right), and MEC (right). Predictions are shown for one (top) or three (bottom, either wells 3, 4, and 5 or wells 6, 7, and 8) well holdouts. Each gray curve represents the median predictions from a single session, with the medians of all sessions denoted by colored diamonds. (**L)** Root mean squared errors of well-identity predictions for the two strategies in **K**, holdout of either single (top) or three (bottom) wells. Horizontal line represents the chance level obtained by shuffling target-well identities during training of the GPR decoder. Filled colored and black diamonds represent individual sessions and their means.

We next focused on the population-level analysis of neural activity manifolds formed by subpopulations of neurons exhibiting significant well-specific firing (total of 4166 out of 9731, 42.81% of OFC neurons; 1504 out of 2684, 56.04% of HPC neurons; 1051 out of 1416, 72.44% of MEC neurons; cells selected as p < 0.05 by ANOVA on single trial neural contributions obtained using singular value decomposition, SVD, see Figure S3 and Methods). To visualize well-specific neural representations in low-dimensional space, we first performed principal component analysis (PCA) on the activity of simultaneously recorded neural ensembles during the well-approach and licking phases.

Figure 2B shows the OFC neural activity from an example session projected on the first three principal components (PCs). The PC projections highlight three properties of the OFC spatial map: 1) trials targeting the same wells are clustered together, signifying robust encoding of individual locations, 2) trials targeting neighboring locations are represented by similar neural ensemble activity, thereby preserving the topological order of well positions along the linear maze, and 3) well locations are mapped on a curved surface such that the representations of track-end wells are brought closer, resulting in a U-shaped manifold. Averaging the PC projections by well identity further revealed a mapping scheme where the distances between neural representations of adjacent wells appear consistent, suggesting that OFC neurons may preserve not only the topological order but also the relative distances between wells, albeit along a curved manifold.

To confirm this intuition, we subsequently assessed the distances between neural representations of individual wells across trials. However, the distance measurement requires representing neural dynamics during the well-approach and licking as single vectors rather than vector sequences. To meet this requirement, we applied tensor component analysis^26^ (TCA) to yield a matrix of trial factors where each row corresponds to an individual trial (referred to as ‘trial factors’) and each column represents its features (referred to as ‘trial components’). These trial components can serve as a vectorized summary of neural population activity dynamics in each trial, capturing its key features while factoring out its temporal dimension. For computing distances, we aimed to focus on a small number of trial components to keep the comparisons within low-dimensional space, as distance measures tend to lose discriminability in higher-dimensional spaces – a manifestation of the phenomenon known as the curse of dimensionality. The number of necessary trial components was estimated based on the decoding performance of well identities by varying the number of trial components. In all three brain regions, a large portion of decoding performance was achieved using only the first few trial components (Figure 2G; n = 8 for OFC, 6 for HPC, and 6 for MEC trial components explaining at least 85% of the saturating decoding probability). This low-dimensional representation is also visually confirmed – plotting the first three trial components (Figure 2D) produced a topology-preserved spatial map at the single-trial level, closely resembling the PC projections of the same data (Figure 2B; see Figure S4 for examples of topology-preserved mapping from all animals used). To estimate the distances of target well representations along the curved manifold, we computed pairwise geodesic distances (see Methods) between trial factors of individual target wells. This revealed that neural activity distance increased with physical distance between the representing wells (Figure 2E). Furthermore, we found a linear relationship between the median neural geodesic distances and the physical distances separating corresponding well pairs in the linear maze (Figures 2F and 2J left). These results confirm that the OFC spatial map preserves both the topological order and the relative distance between well locations.

Similarly, we also examined the spatial mapping of target wells by neural ensembles in HPC and MEC. PCA and TCA-based dimensionality reduction and visualization revealed that both HPC and MEC ensembles also form topology-preserved maps of well locations, which appear to reside on a curved surface (Figures 2H-2I). Furthermore, the geodesic distance analyses confirmed that neural representational distances scale linearly with the physical distances between corresponding wells (Figure 2J: mean ± s.e.m. of correlation coefficients between neural and physical distances: 0.62 ± 0.025, n = 62 sessions in OFC; 0.57 ± 0.042, n = 37 sessions in HPC; 0.68 ± 0.048, n = 21 sessions in MEC). In contrast, computing Euclidean distances, instead of geodesic, severely impaired these linear relationships in all three brain regions, supporting the idea that these maps reside on curved manifolds (Figure S4).

Finally, we tested if the OFC neurons encode well locations as a continuum such that the representation of each well is defined by those of its neighboring wells. We used GPR to predict the location of a given well based on the model trained on the neural representations of the remaining wells in the low dimensional TC space. We found that the identity of the well held out from the training could be predicted significantly better than chance levels (Figures 2K and L, top panels; mean ± s.e.m. of RMSEs: 1.31 ± 0.024, chance level 1.93, n = 62 sessions). To further test the uniformity of spatial coding over a larger scale, we performed a more stringent GPR analysis in which three wells, either from 3 to 5 or from 6 to 8, were held out from the training, and we still found that the positions of held-out wells could be regressed from the representations of the remaining ones (Figures 2K and L, bottom panels; mean ± s.e.m. of RMSEs: 1.62 ± 0.043, chance level 2.47, n = 51 sessions). Importantly, these GPR analyses treated reward locations as continuous rather than discrete variables, and the successful decoding of well identity highlights the OFC’s capacity to represent well locations along a spatial continuum. We further confirmed successful regression of held-out wells from TC representations of both HPC and MEC neural ensembles (Figures 2K and L; mean ± s.e.m. of RMSEs: 1.1 ± 0.045, chance 1.87, n = 37 sessions for 1-well hold-out; 1.6 ± 0.067, chance 2.46, n = 28 sessions for 3-well hold-out in HPC; 1.04 ± 0.033, chance 1.91, n = 21 sessions for 1-well hold-out; 1.4 ± 0.075, chance 2.5, n = 15 sessions for 3-well holdout in MEC), which is consistent with the continuous spatial activity manifolds observed in both regions (Figures 1B and C). The results of both the geodesic distance and the GPR-based analyses were largely robust to the number of TC dimensions, although distance discrimination declined at higher dimensions (Figure S4).

Taken together, these results point to a novel spatial mapping scheme in the OFC that preserves the topological arrangements of locations in an environment.

### The OFC spatial map scales with distance

The fact that the brain’s spatial maps in OFC, HPC, and MEC preserve relative distances between well positions suggests that these maps may encode not only the topological arrangements of positions but also the metric distances between them. However, it is also possible that these representations may simply reflect the sequential order of well positions along the maze without explicitly encoding the physical distances. To distinguish these possibilities, in a subset of sessions, we omitted an intermediate well, ensuring that the animals did not target this well as a navigational goal. We found that the geodesic distance between neural representations of two target wells separated by an unused well (i.e., two wells apart in physical space) was greater than the distances between two adjacent used wells (Figure S5; mean ± s.e.m. of geodesic distance (AU), 0.432 ± 0.018 for adjacent used wells, 0.496 ± 0.029 for wells separated by an unused well, p = 0.016 by sign rank test, n = 14 sessions), indicating that the OFC spatial map reflects actual physical separation rather than the ordinal position.

Next, we asked whether the OFC map possesses a uniform distance metric that is preserved across different navigation contexts. Such a common metric would allow the map to scale with the size of the environment, which is advantageous for the appropriate planning of navigational strategies adapted to each environment. To test this possibility, rats were trained to navigate not only the standard 2-meter maze, but also perform the same task in a 4-meter linear track with ten equally spaced reward wells (Figure 3A). The rats performed goal-directed navigation under the same task rule, once in the standard linear maze (“linear”) and once in the large maze (“large”; n = 10 sessions, 5 rats for OFC; n = 7 sessions, 3 rats for HPC; from n = 7 sessions, 2 rats for MEC). We observed that OFC neurons exhibit a larger range of firing rates to represent individual wells in the large maze compared to the linear maze (Figure 3B). A similar increase in firing-rate range was also visible in MEC neurons but not in HPC neurons (Figure 3B).

**Figure 3:**
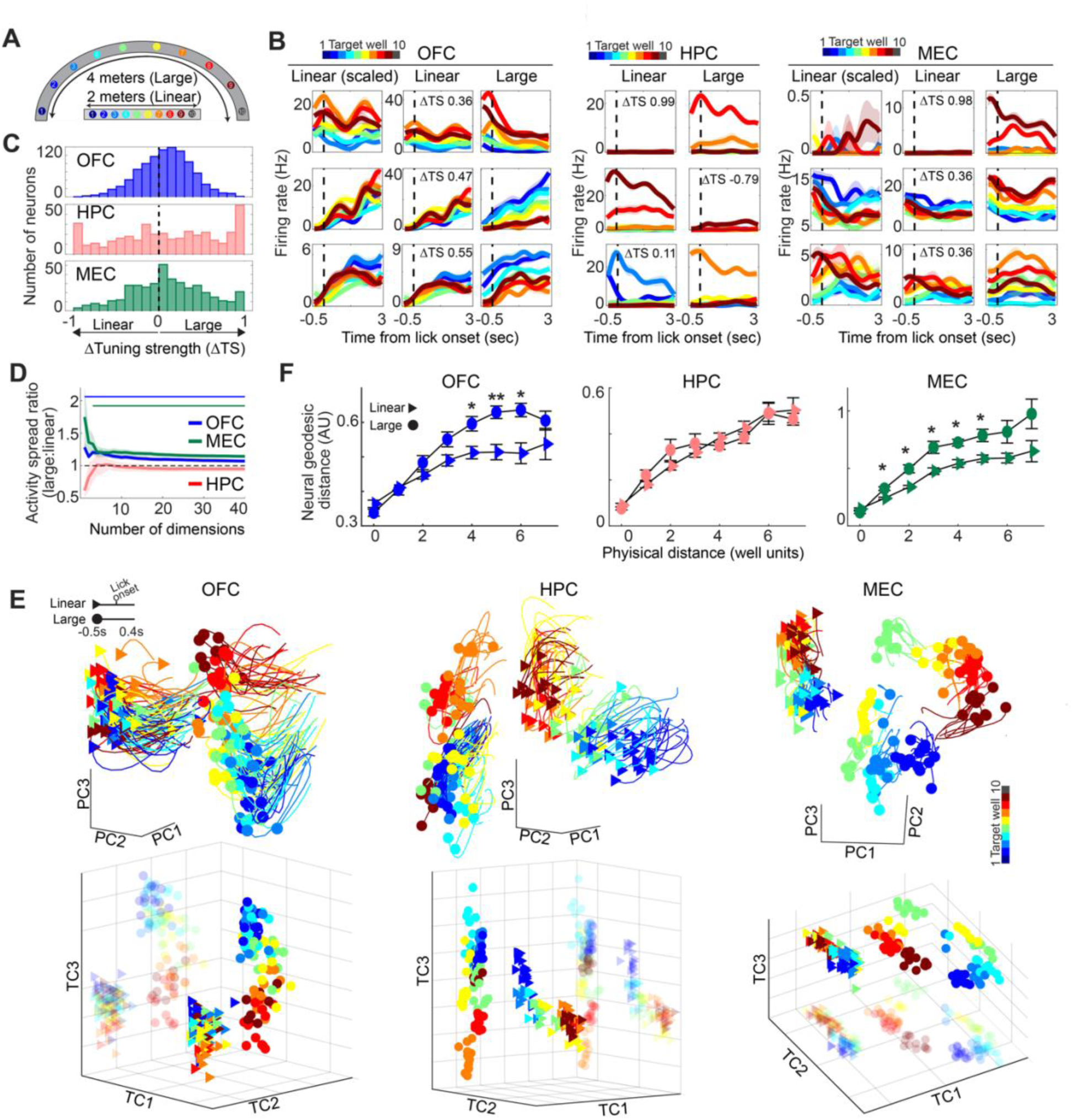
Spatial maps in OFC and MEC, but not HPC, scale with the maze size. **(A)** Schematic of the linear and the large mazes. **(B)** Firing rates of three representative neurons, each from OFC (left), HPC (middle), and MEC (right). For OFC and MEC, the middle and right panels show firing rates in the linear and the large mazes using the same scales, while the linear-maze firing rates in the left panels are rescaled to reveal target-well specific activity. The tuning strength differences (ΔTS) are shown for each neuron. **(C)** Histogram of ΔTS (see main text) of individual neurons in OFC (top), HPC (middle), and MEC (bottom). **(D)** Ratios of neural activity spread (large to linear) at lick onset for OFC, HPC, and MEC neural ensembles, plotted as a function of the number of principal components. Horizontal bars on top indicate ranges where the ratio significantly differs from 1 (p < 0.05 in Wilcoxon signed-rank test). Shown are means (solid) ± s.e.m. (shading). **(E)** Ensemble neural activity of OFC (left), HPC (middle), and MEC (right) in the linear (triangular head) and the large (circular head) mazes from an example paired session. Top panel plots neural activity projected on three principal components. Neural activity from 500 ms before to 400 ms after lick onset is shown with a color-code based on the target-well identity (each thin line). Bottom panel shows the first three TCA trial components of the same sessions. Translucent triangles and circles are projections on a single plane, highlighting the enlarged mapping of the large maze in OFC and MEC, but not HPC. To facilitate comparisons of neural representations between the two mazes, the representations from one maze are shifted along a single dimension. **(F)** Geodesic distances between neural representations of well pairs in OFC (left), HPC (middle), and MEC (right), plotted against physical distances (in well units) for the linear and the large mazes. Note that the well unit (separation) in the large maze is twice as large as that in the linear maze. Shown are means (filled triangle and circle) ± s.e.m. (errorbars). *p < 0.05, **p < 0.01 by Wilcoxon sign-rank test.

To quantify the range of target-well coding, we regressed the trial-wise contributions of individual neurons to the corresponding well-identities in a given maze, derived from singular value decomposition (SVD). This analysis resulted in a vector of regression coefficients for individual wells in each maze, and the magnitude of these vectors was defined as the neuron’s tuning strength for the corresponding mazes (see Figure S3 and Methods). The tuning strengths of OFC neurons were higher in the large maze compared to the linear maze, as evident from an overall shift in the distribution of normalized tuning strength difference (ΔTS), a pattern also observed in the MEC neuron population (Figure 3C; median ΔTS: 0.092, n= 890 neurons in OFC, 0.109, n= 381 neurons in HPC, and 0.089, n= 405 neurons in MEC). In contrast, HPC neurons did not exhibit a similar distribution of ΔTS, but 10.5% (40/351 neurons with ΔTS <-0.8) and 18.6% (71/361 neurons with ΔTS > 0.8) of HPC neurons exhibited place-specific firing only in the linear or large maze, respectively, consistent with global remapping^9^. Maze-specific firing was also observed in 9.8% of MEC neurons but was almost absent in OFC neurons (9/405 with ΔTS <-0.8 and 31/405 neurons with ΔTS > 0.8 in MEC; 2/890 with ΔTS <-0.8 and 9/890 neurons with ΔTS > 0.8 in OFC). Taken together, individual OFC and MEC neurons, but not HPC neurons, represent well locations in the large maze with a greater range of firing rates compared to those in the linear maze.

We next asked if this increased range of firing rates of individual neurons contributes to an expanded representation of well positions in the large maze at the neural population level. To test this idea, we first confirmed whether well locations in the large maze are topologically represented, as in the standard linear maze. We identified distinct neural representations of individual well positions in the large maze, leading to significant decoding of well identity (Figure S5). In addition, visualization of OFC neural activity in both PC and TC spaces revealed topology-preserved well representations, and furthermore, the geodesic distances between neural well representations scaled linearly with the physical distances between wells. GPR analysis further confirmed that the position of a given well could be predicted from the neural representations of the remaining wells (Figure S5). Topology-preserved mapping of well positions was also observed in the MEC neural ensembles. However, despite highly well-specific representations, HPC neural ensembles did not exhibit a clear topological ordering of wells in the large maze (Figure S5).

We then compared the scale of spatial maps between the two mazes. Because the same sets of neurons were recorded in both mazes during paired sessions, we were able to directly compare the spatial maps for the two mazes at the single-trial level. We first compared the spread of neural activity, defined as the mean-centered distances of the population activity at lick onset, averaged across trials. To account for dimensionality effects on distance measurements, we computed activity spreads across increasing numbers of PC dimensions used for neural activity projections. In both OFC and MEC neural ensembles, activity spreads were greater in the large maze than in the linear maze across the first 40 PC dimensions. In contrast, HPC ensembles showed no significant difference in activity spread between the two mazes (Figure 3D; the ratio of activity spread in large to linear track differed from 1 at p < 0.05 for dimensions 1 to 40 for OFC; for dimensions 3 to 40 for MEC; and p > 0.05 at all tested dimensions for HPC; p-value obtained by sign rank test).

In support of the increased activity spread, visualizing neural representations at the single-trial level in both PC and TC spaces revealed expanded spatial mapping of well positions in the large maze for OFC and MEC ensembles, but not for HPC (Figure 3E). We subsequently computed the geodesic distance between neural representations of well pairs in both mazes using a small number of TC dimensions (n=11, 6, and 8 dimensions in OFC, HPC, and MEC, respectively; dimensions were selected such that they explain 82.77%, 85.43%, and 87.88% of the peak target well decoding averaged over both linear and large mazes; see Figure S5 and Methods for details). Neural geodesic distances corresponding to the same number of well separations were significantly greater in the large maze compared to the standard linear maze for both OFC and MEC neural ensembles, indicating a scalable spatial representation depending on the maze size. In contrast, no such scaling was observed in HPC neural ensembles (Figure 3F). We further confirmed that this increase in geodesic distances in the large maze was robust to variations in the number of TC dimensions (Figure S5).

Our results demonstrate that the OFC spatial map scales with the distance between encoded positions, both at the level of individual neurons and neural ensembles. This scaling property was also observed in MEC, but was absent in HPC, consistent with previous reports that MEC uniquely encodes a distance metric by integrating locomotion signals^8,27,28^. Notably, the presence of distance-dependent scaling in the OFC map suggests that the OFC, too, contains a distance metric - a feature absent in HPC.

### The OFC spatial map generalizes across environmental contexts

Multiple lines of behavioral evidence suggest that animals have the ability to transfer learned behaviors in one environment to another distinct but similar one, suggesting the existence of a schematic map that generalizes across environments^29,30,14,16,1^. However, place cells in the hippocampus drastically change their firing patterns between different environmental contexts, resulting in orthogonal spatial maps^10,9^ (termed ‘global remapping’). Similarly, spatial representations of grid cells in MEC also orthogonalize when environmental cues are changed in the same maze^11^. These results suggest that the maps in HPC and MEC are unlikely to support the generalization of navigation behaviors across environments. We thus tested whether the OFC spatial map might instead generalize the arrangement of encoded spatial positions across environments.

We conducted paired sessions where the same task was carried out on identical linear mazes located in two different rooms while recording from the same neurons (n = 11 paired sessions from 5 rats for OFC; n = 8 paired sessions from 3 rats for HPC, n = 6 paired sessions from 2 rats for MEC). HPC and MEC neurons exhibited prominent changes in spatial firing patterns, with some neurons becoming silent in one of the rooms (Figure 4A). In contrast, most OFC neurons were active in both rooms and maintained similar firing patterns at individual wells (Figure 4A). To quantify the change in tuning properties for each neuron, we computed the ΔTS and the tuning distance, defined as the cosine distance between the regression coefficient vectors in the two rooms (as described in the previous section). Plotting both parameters for all neurons revealed distinct patterns of changes across the three brain regions. HPC neurons were largely divided into two populations, each active in one of the two rooms, with overall high tuning distances (30.72%, 133/433 neurons with |ΔTS| > 0.8, median tuning distance 1.38). In contrast, the ΔTS of OFC neurons were mostly clustered around 0 with broadly distributed tuning distances (0.51%, 5/973 neurons with |ΔTS| > 0.8, median tuning distance 1.13). MEC neurons showed an intermediate pattern with broadly distributed ΔTS along with an overall high tuning distance (10.91%, 37/339 neurons with |ΔTS| > 0.8, median tuning distance 1.46). To assess remapping at the population level, we computed pairwise cosine distances between population activity vectors representing individual trials (see Methods). Example distance matrices from one paired session for each brain region are shown in Figure 4C. Focusing on the cosine distances between the two rooms (top-right quadrant of the distance matrix) revealed that the OFC neural ensemble maintained a similar pattern of representational distances between target wells both within and across rooms. In contrast, neural ensembles in HPC and MEC exhibit uniformly high representational distances between target wells across the two rooms (Figures 4C and S6), consistent with the previously reported orthogonalization of spatial maps in two distinct environments^9,11^. To quantify this observation, we measured the difference in the representational distances between well pairs across and within rooms (normalized to their sum) and found significantly lower values in OFC compared to HPC and MEC, suggesting greater similarity in well-position mapping between the two rooms in OFC (Figure 4D; mean ± s.e.m. of normalized differences: 0.074 ± 0.004, 0.163 ± 0.014, and 0.233 ± 0.034 for OFC, HPC, and MEC respectively; p = 0.004 OFC vs HPC; p = 0.0002 OFC vs MEC; and p = 0.559 HPC vs MEC by Kruskal-Wallis test followed by pairwise comparison). Overall, the above analyses point to a unique feature of the OFC map in maintaining spatial representations across environments, unlike those in HPC and MEC.

**Figure 4:**
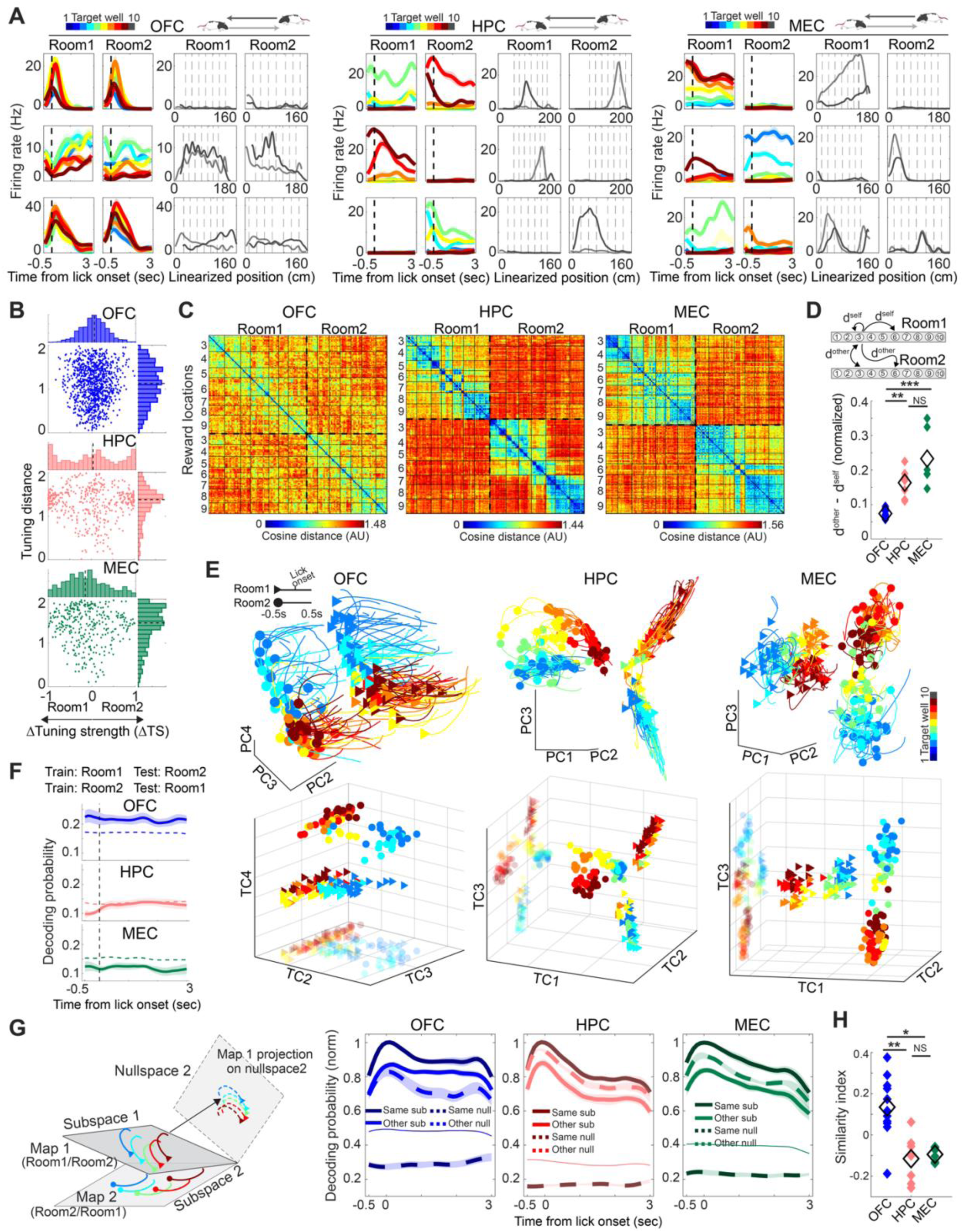
OFC maps uniquely preserve topological structure across environments. **(A)** Firing rates of three representative neurons from OFC (left), HPC (middle), and MEC (right) on the linear mazes located in two different rooms. For each neuron, the two left panels show target-well specific firing rates aligned to lick onset, and the right two panels plot maze-position-dependent rates in the two rooms. Activity for the two running directions is plotted with separate shading (light and dark grey). Dotted vertical lines represent well locations (wells 3-9 were used). (**B)** Tuning distance versus ΔTS plotted for OFC (top), HPC (middle), and MEC (bottom) neurons. Each dot corresponds to a single neuron. Marginal distributions are plotted on the top and right of each panel. Dotted lines represent the medians. **(C)** Representational distance matrices between target-wells across the two rooms, for example sessions from OFC (left), HPC (middle), and MEC (right). Each pixel depicts the cosine distance between population vectors of neural activity for a pair of trials ( methods). Trials are grouped according to the target well identity, and trials from the two rooms are separated by thick dotted lines. **(D)** Top: schematic describing the metrics d^self^ and d^other^, defined as neural activity distances between trials targeting paired wells within and across rooms, respectively. Bottom: the difference between d^other^ and d^self^ normalized to its sum and averaged across all well pairs in a session. Filled colored and open black diamonds represent individual sessions and their means, respectively. **(E)** Ensemble neural activity on the linear mazes in the two rooms from a representative paired session, each from OFC (left), HPC (middle), and MEC (right). Top panel shows single-trial neural activity, color-coded by the target well identity, and projected on three principal components. Bottom panels show trial factors in the corresponding sessions projected on three TCA trial components. Translucent triangles and circles are projections of the trial factors on a single plane, highlighting the similarity in OFC spatial representations between the two rooms, in contrast to largely orthogonal arrangements in those of HPC and MEC. To facilitate comparisons of neural representations between the two rooms, the representations from one room are shifted along a single dimension. **(F)** Probabilities of target-well decoding in one room using a decoder trained on the neural activity in the other room. Thin dotted lines show the chance levels. Shown are means (solid) ± s.e.m. (shading). **(G)** Left: schematic of the subspace-based decoding analysis. Right: probabilities of target-well decoding from neural activity in OFC, HPC, and MEC, projected on the distinct subspaces occupied by the spatial maps in the two rooms, and their corresponding null spaces ( methods). Shown are means (solid and thick dotted lines) ± s.e.m. (shading). Thin lines show the chance levels of decoding on the same subspace. For each paired session of two-room recordings, the decoding probabilities are normalized to those on the same subspace at lick onset. **(H)** Subspace similarity indices (see main text), plotted for each paired session (individual colored diamonds) from each of the three brain areas. Open black diamonds depict the means. *p <0.05, **p <0.01, ***p < 0.001 based on Kruskal-Wallis test followed by post hoc pairwise comparison.

We next explored how this unique feature of the OFC map is supported by the neural representations at the population level. In HPC, global remapping is thought to arise when place cells either shift their firing fields to entirely different locations in a new environment or fire selectively in only one of the environments. At the ensemble level, such remapping results in spatial maps for individual contexts occupying orthogonal subspaces (where subspace refers to a subset of axes that contain the map, Figure S6). However, a ‘milder’ form of remapping is also possible in which the two maps reside in similar subspaces but differ in the orientation of their location representations (Figure S6). These two modes of remapping have distinct functional implications: changes within a shared subspace can support behavioral adaptations across environments, while out-of-subspace remapping will hinder generalization and necessitate re-learning in a new context^31^. The high cosine distances between HPC and MEC population vectors across rooms suggest that the spatial maps in these brain regions occupy highly dissimilar subspaces. In contrast, the lower cosine distances of the OFC map across trials from different rooms suggest that the OFC map resides in similar subspaces and maintains a largely consistent spatial coding scheme across contexts.

To confirm these predictions, we examine how the two spatial maps in each brain region are oriented relative to each other. PCA and TCA-based visualizations of OFC neural ensemble activity from two rooms in paired recording sessions revealed that target-well representations along the maze positions appear aligned, and importantly, the two maps occupy similar subspaces (Figure 4E). In contrast, spatial maps in HPC and MEC from the two rooms reside along orthogonal axes, exhibiting a cross-shaped structure in the TC space (Figure 4E bottom). To further quantify whether the two spatial maps share a consistent position coding scheme, we trained a decoder on well identity in one room and tested its performance on the other. The decoder successfully predicted well identity on the held-out maze above chance levels in the OFC but not HPC and MEC neural ensembles (Figure 4F; mean ± s.e.m. decoding probability at lick onset: 0.219 ± 0.021 in OFC; 0.113 ± 0.009 in HPC; 0.115 ± 0.011 in MEC), supporting the idea that the OFC map generalizes spatial information across environments. To further characterize representational alignment across environments, we computed the LDA axis that maximally separated neural representations of well identity in each room. The angle between the two LDA axes was significantly smaller in OFC than in HPC and MEC (Figure S6), indicating that OFC maintains consistent representations across environments. Together, the decoding and LDA results suggest that the OFC employs a shared spatial coding scheme across environments, in contrast to the remapped representations observed in HPC and MEC.

Finally, to test how remapping might affect a downstream brain region that is poised to read out neural activity from a given subspace^32^, we asked how much spatial information is shared between the representational subspaces of individual environments. Specifically, we quantified how much spatial information in one room is preserved in the neural subspace in another room and how much is excluded in the corresponding nullspace orthogonal to that subspace. For fair comparison, we sought to equalize the dimensionality of the subspaces and their corresponding nullspaces. Specifically, we started by operationally defining null space as a set of axes (PC dimensions) that explain the last 10% variance in neural activity during approach and licking of target wells within a given maze, while representational subspace comprised an equal number of leading PC dimensions that explained a median of 87.21% variance across all three brain regions (see Methods). We then evaluated how much spatial information can be transferred by projecting neural activity in one room onto the representational subspace defined in the other (‘other’s subspace’) and onto the corresponding nullspace (‘other’s nullspace’) by comparing the decoding performances between these two projections (Figure 4G left). If the two maps reside on similar representational subspaces, decoding performance of well identity in ‘other’s subspace’ would exceed that in ‘other’s nullspace’ because the other’s subspace would provide a geometrical structure that allows separation of well positions similar to the original subspace. Consistent with this prediction, OFC ensembles show significantly higher decoding performance in the ‘other’s subspace’ than in the ‘other’s nullspace’ (Figure 4G). In contrast, HPC and MEC exhibited an opposite trend, with better decoding in the nullspaces. To quantify this effect, we defined the subspace similarity index as the difference in decoding performance between ‘other’s subspace’ and ‘other’s nullspace’, normalized to the corresponding decoding difference in the original environment between ‘same subspace’ and ‘same nullspace’. The index was significantly higher in OFC compared to HPC and MEC (Figure 4H; mean ± s.e.m.: 0.13 ± 0.046,-0.11 ± 0.038, and-0.09 ± 0.014 for OFC, HPC, and MEC respectively; p = 0.006 OFC vs HPC; p = 0.015 OFC vs MEC; and p = 0.999 HPC vs MEC by Kruskal-Wallis test followed by pairwise comparison), confirming that the OFC maps reside in shared subspaces between environments. Importantly, this subspace similarity in OFC is not an artifact of similar temporal dynamics of individual neurons in the two different rooms (Figure 4A), as cross-environment similarity in spatial mapping was observed even when temporal dynamics were taken out, as shown in the population vector analysis (Figures 4C-D) and in the TCA-based visualization (Figure 4E).

Despite these differences in neural subspace sharing across environments in different brain regions, the decoding performance for well identity within each environment was comparable across OFC, HPC, and MEC (Figure S6). In addition, using the same geodesic distance analysis as in Figure 3F, we confirmed that the scales of the spatial maps formed for identical mazes in the two rooms were comparable across all regions (Figure S6). These results suggest that spatial maps were properly formed in both rooms, although the relative orientations of their maps differed across brain regions. Further, we verified that the pronounced spatial remapping observed in HPC and MEC was not due to instability of neural representations because spatial representations remained stable between paired sessions on the same maze in the same room, both at the single neuron and the neural ensemble levels in all three brain regions (Figure S7).

Taken together, these findings suggest that the OFC forms a schematic spatial map that generalizes across environments, which contrasts strikingly with the maps in HPC and MEC that exhibit pronounced remapping and context specificity.

### The OFC preserves map structure across maze geometries

Global remapping of the HPC spatial map is known to be triggered not only by changes in rooms but also by alterations in maze geometry, even when the underlying task structure remains unchanged^9^. To test if the OFC spatial map generalizes across maze geometries, we introduced a circular maze with a circumference of 2 meters, ten equally spaced reward wells, and a barrier between wells 1 and 10, making it topologically and metrically identical to the original linear maze (Figure 5A). Rats rapidly adapted to the new geometry and performed paired sessions on the linear and the circular mazes with similar accuracy (Figure S8; n = 12 paired sessions from 5 rats for OFC; n = 9 paired sessions from 3 rats for HPC, n = 6 paired sessions from 2 rats for MEC). As in the linear maze, OFC neurons in the circular maze exhibited target-well specific firing, often preserving the order of the well locations in their firing pattern (Figure 5B). In contrast, HPC neurons tended to fire at single wells, and MEC neurons showed a mixture of both broad and narrow tunings to target wells (Figures 5B and S8). Decoding analyses confirmed that target-well identities in the circular maze could be accurately predicted from neural ensemble activity in all three brain regions, suggesting the formation of distinct well-position representations (Figure S8).

**Figure 5:**
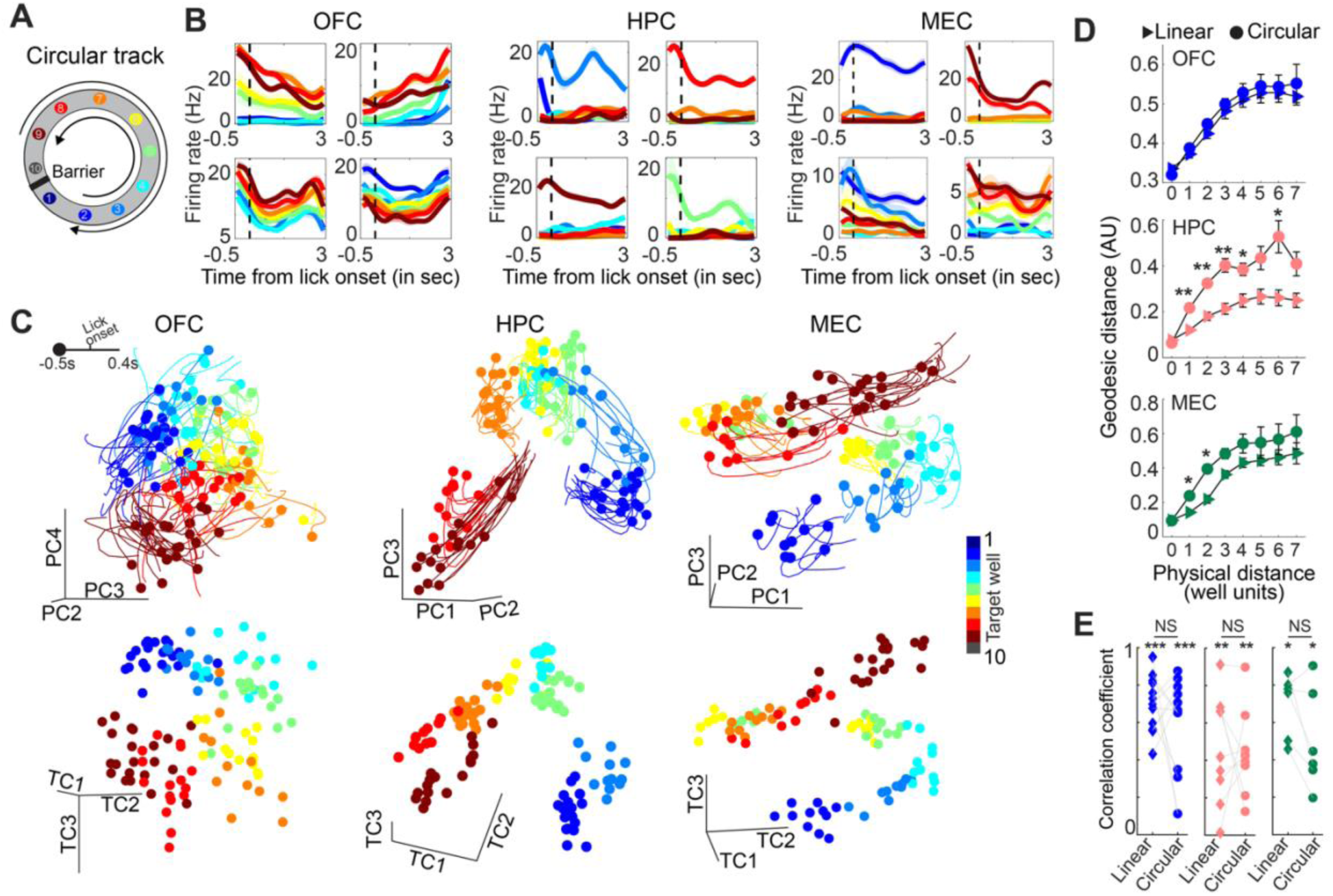
Spatial maps in OFC, HPC, and MEC preserve topology along the circumference of a circular maze. **(A)** Schematic of the circular maze. **(B)** Firing rates of four representative neurons each from OFC (left), HPC (middle), and MEC (right). **(C)** Ensemble activity of neurons in OFC (left), HPC (middle), and MEC (right) on the circular maze from one example session each. Top panels plot single-trial neural activity, color-coded by target well identity, and projected on three principal components. Bottom panels show the first three TCA trial components of the corresponding session. Note the preservation of the relative well locations in the OFC spatial map. **(D)** Geodesic distance between well representations in the TC space plotted against the physical distance between corresponding wells for both linear (diamond) and circular (circle) mazes. Shown are means ± s.e.m. across sessions (same convention as in Figures 2J and 3F). **(E)** Correlation coefficients between geodesic neural-representation distances and physical distances. Diamond and circle connected by a line represent the correlation coefficients in the linear and circular mazes from a paired session with the same neural population recorded. Correlation coefficients for individual mazes are significantly larger than zero, with no significant differences between mazes across all brain regions. In panels D and E, *p<0.05, **p<0.01, ***p<0.001 by Wilcoxon signed-rank test.

Visualizing the spatial mapping on the circular maze using dimensionality reduction revealed that OFC neurons preserved the topological arrangement of well positions during approach and licking (Figure 5C). To quantify this feature, we measured the geodesic distances between trial factors derived from TCA and found that the neural distances in OFC scaled linearly with the physical distances between corresponding wells in both the circular and linear mazes (Figures 5D and 5E; mean ± s.e.m. of correlation coefficients in linear and circular mazes: 0.703 ± 0.043 and 0.567 ± 0.079). GPR-based analyses also confirmed that the well identities, held out from the training set, could be successfully predicted from the representations of the remaining wells, further pointing to a topology-preserved map in OFC (Figure S8). Furthermore, neural geodesic distances between pairs of equally separated wells were nearly identical between the two mazes (Figure 5D) despite the different geometries. This result suggests that the OFC map scales according to path length along the circumference, rather than Euclidean distance (i.e., chord lengths), reinforcing our results from the large maze (Figure 3).

Topology-preserved mapping in the circular maze was also observed in HPC and MEC neural ensembles (Figure 5C). In both regions, neural geodesic distances between pairs of equally separated wells were correlated with their physical path lengths in both mazes, although the spatial map on the circular maze appeared more expanded in the HPC (Figures 5D and 5E; mean ± s.e.m. of correlation coefficients in linear and circular mazes: 0.462 ± 0.096 and 0.439 ± 0.075 in HPC; 0.694 ± 0.07and 0.504 ± 0.109 in MEC). The increased scale of the HPC map could be explained by enhanced tuning strength (ΔTS) of individual HPC neurons in the circular maze, while OFC and MEC neurons exhibited only modest changes between the two mazes (Figure 6B; median ΔTS: 0.062 in OFC, 0.23 in HPC, and 0.108 in MEC).

**Figure 6:**
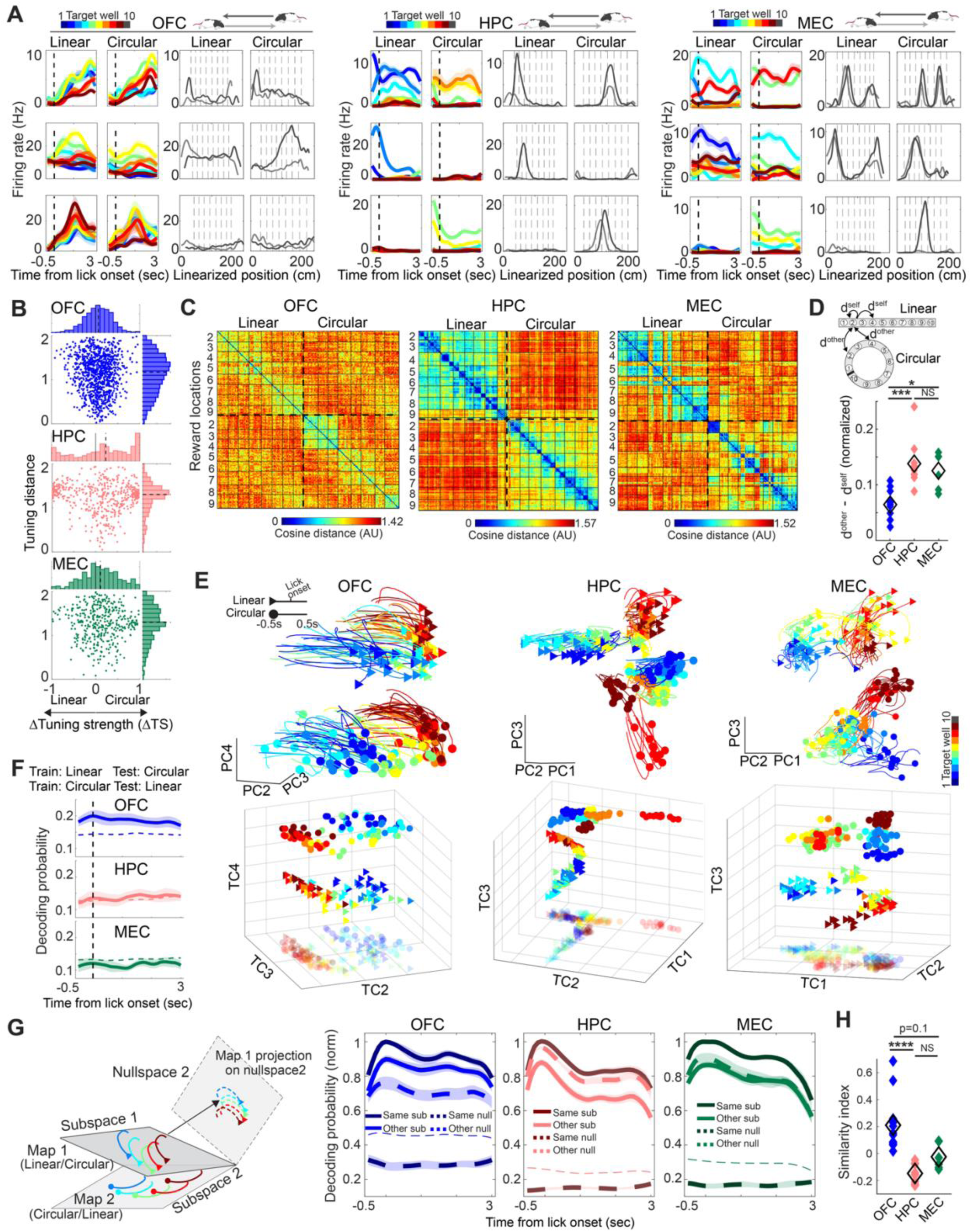
OFC spatial maps uniquely preserve topological structures across maze geometries. **(A)** Firing rate of three representative neurons each from OFC (left), HPC (middle), and MEC (right) in the linear and circular mazes. For each neuron, the two left panels show target-well specific firing rates aligned to lick onset, and the right two panels plot firing rates along the positions in the two mazes. Activity for the two running directions is plotted in separate shades (light and dark grey). Dotted vertical lines represent well locations (wells 2-9 were used). **(B)** Tuning distance versus ΔTS plotted for OFC (top), HPC (middle), and MEC (bottom) neurons. Each dot corresponds to a single neuron. Marginal distributions are plotted on the top and right of each panel. Dotted lines represent medians. **(C)** Representational distance matrices between target-wells across the two mazes, for example paired sessions from OFC (left), HPC (middle), and MEC (right) neural ensembles. Each pixel depicts the cosine distance between the population vectors of neural activity for a pair of trials. Trials are grouped according to the target well identity, and trials from the two mazes are separated by thick dotted lines. **(D)** (top) Schematic describing the metrics d^self^ and d^other^, which denote representational distances between well pairs within and across mazes, respectively. (bottom) The difference between d^other^ and d^self^, normalized to its sum and averaged across all well-pairs in a session. Filled colored and open black diamonds represent individual sessions and their means, respectively. **(E)** Ensemble neural activity in the two mazes from representative paired sessions recorded from OFC (left), HPC (middle), and MEC (right). Top panel shows single-trial neural activity, color-coded by the target well identity and projected on three principal components. Bottom panels show trial factors of the corresponding sessions projected on three trial components. Translucent triangles and circles are projections of the trial factors on a single plane. To facilitate comparisons of neural representations between the two mazes, the representations from one maze are shifted along a single dimension. **(F)** Decoding probabilities of target wells in a given maze using a decoder trained on the other maze. Thin dotted line shows the chance level. Shown are means (solid) ± s.e.m. (shading). **(G)** Left: schematic of the subspace-based decoding analysis. Right: Decoding probabilities of target wells from neural activity projected on the distinct subspaces occupied by the spatial maps in the two mazes, as well as their corresponding null spaces. Shown are means (solid and thick dotted lines) ± s.e.m. (shading). Thin lines show the chance levels for decoding on the same subspace. For each paired session, the decoding probabilities are normalized to those on the same subspace at lick onset. **(H)** Subspace similarity indices (see main text), plotted for each paired session (individual colored diamonds) from all three brain areas. Open black diamonds depict the means. *p < 0.05, ***p < 0.001, ****p < 0.0001 based on Kruskal-Wallis test followed by post hoc pairwise comparison.

Next, we focused on the relationship between spatial maps in the linear and circular mazes and examined the degrees of remapping across the three brain regions. Representative neurons shown in Figure 6A highlight that individual OFC neurons fired in both mazes and exhibited partly similar tuning to individual wells. In contrast, HPC and MEC neurons either fired at different locations between mazes or became silent in one of the mazes, consistent with the features of global remapping. Quantifying the ΔTS and tuning distance revealed that HPC neural population was largely split into two groups, each preferring one of the mazes (162/485 neurons, 33.4% with |ΔTS|>0.8), along with overall high and narrowly distributed tuning distances (median: 1.3). In contrast, the OFC neural population maintained consistent tuning strengths across mazes (11/1055 neurons, 1.04% with |ΔTS|>0.8) with broadly distributed tuning distances (median: 1.17). MEC neurons exhibited an intermediate pattern with modest preference for one maze (35/396 neurons, 8.84% with |ΔTS|>0.8) but overall high tuning distances (median: 1.29). Comparing well-position representations between paired sessions using population vector analysis revealed that OFC neurons encode well pairs by preserving similar representational distances across the two mazes (Figures 6C-6D). This similarity was absent in HPC and MEC neural ensembles, with HPC population vectors maintaining uniformly high representational distances between cross-maze well pairs (Figures 6C-6D and S8). These results suggest that the spatial maps in OFC, and to a lesser extent MEC, occupy similar subspaces between mazes with different geometries, but not those in HPC.

However, important differences exist between the spatial maps in OFC and MEC. Visualizing the spatial representations from both linear and circular mazes from paired sessions using dimensionality reduction revealed that OFC spatial maps maintained similar orientation across the two geometries, whereas the MEC map did not (Figure 6E). To quantify orientation similarity, we trained a decoder for well identity on one maze and tested it on the other. Decoding performance was significantly above chance in OFC but not in HPC and MEC (Figure 6F; mean ± s.e.m. of decoding probabilities at lick onset: 0.199 ± 0.017 in OFC; 0.135 ± 0.022 in HPC; 0.121 ± 0.015 in MEC). We finally examined whether the spatial maps from the two mazes occupied similar subspaces using the subspace-and nullspace-based decoding analysis described in Figure 4. In OFC, projecting neural activity from one maze on ‘other’s subspace’ yielded higher decoding performance than projecting on ‘other’s nullspace’ (Figures 6G-H), confirming shared map subspaces across maze geometries. In contrast, HPC maps showed the opposite trend, with decoding performance higher in ‘other’s nullspace’ than in ‘other’s subspace’, indicating highly dissimilar spatial representations between the mazes. MEC maps showed similar decoding performance between ‘other’s subspace’ and ‘other’s nullspace’ (mean ± s.e.m. of similarity indices: 0.21 ± 0.059 in OFC,-0.147 ± 0.023 in HPC,-0.024 ± 0.039 in MEC). Hence, these results reveal that changes in maze geometry led to different levels of remapping across brain regions: OFC spatial maps preserve subspaces and orientations; MEC maps share similar subspaces but undergo reorientations; and HPC maps reside in highly dissimilar subspaces partly due to a large fraction of HPC neurons that fire selectively in only one of the two mazes.

Taken together, these findings suggest that, as long as the relative topological arrangement of locations is preserved, OFC neurons form a topology-preserved schematic map that generalizes across rooms (Figure 4) and maze geometries (Figure 6).

### OFC map formation occurs independently of HPC and MEC

Previous studies have suggested that spatial representations in brain regions outside the HPC and MEC depend on inputs from the HPC^33–35^. However, our findings highlight fundamental differences in spatial mapping between the OFC and the canonical spatial areas HPC and MEC. This led us to hypothesize that the OFC map formation may occur independently of inputs from HPC or MEC. To test this hypothesis, we performed lesions of the entire dorsal and ventral HPC (Figure 7A, 2 rats) or the MEC (Figure 7B, 2 rats) in two separate groups of rats. Surprisingly, all lesioned rats learned the goal-directed navigation task with learning curves comparable to those of intact subjects (Figure S9). Once trained, we recorded neural activity from the OFC (Figure S9) as the animals performed the task on the linear maze (n = 11 sessions from HPC lesions, 12 sessions from MEC lesions). OFC neurons in the lesioned animals exhibited target-well-specific firing (908 out of 1609, 56.43% neurons in HPC lesions; 670 out of 1605, 41.74% neurons in MEC lesions; cells selected as p < 0.05 by ANOVA). Moreover, a subset of neurons exhibited firing patterns reflecting the spatial order of well positions (Figure 7C-D), as observed in the intact animals. At the neural ensemble level, well positions were distinctly represented, with decoding performance significantly exceeding chance levels (Figures 7G and S9; mean ± s.e.m. of decoding probabilities at lick onset: 0.452 ± 0.034 for HPC lesions; 0.314 ± 0.018 for MEC lesions). To visualize the neural representation structure, we performed PCA and TCA on the neural activity during approach and licking, revealing topology-preserved maps of well positions in both HPC-and MEC-lesioned animals (Figures 7E-F). Consistent with these observations, we found significant correlations between the geodesic distances of neural representations and the physical distances between corresponding wells (Figure 7H; mean ± s.e.m. of correlation coefficients: 0.53 ± 0.071 for HPC lesions; 0.55 ± 0.056 for MEC lesions). Furthermore, even when one or three wells were held out from the training of decoder for well identify, GPR-based analyses confirmed that held-out well positions could be predicted above chance levels, implying topology preservation of OFC maps across extended spatial scales, independently of HPC or MEC (Figures 7I-J; mean ± s.e.m. of RMSEs: 1.38 ± 0.087, chance 2.05, n = 11 sessions for 1-well hold-out; 1.65± 0.109, chance 2.54, for 3-well holdout in HPC lesions; 1.49 ± 0.057, chance 2.03, n = 12 sessions for 1-well hold-out; 1.71 ± 0.095, chance 2.58, for 3-well hold-out in MEC lesions). Overall, these results confirmed that the OFC can form topology-preserved spatial maps even in the absence of inputs from HPC or MEC.

**Figure 7:**
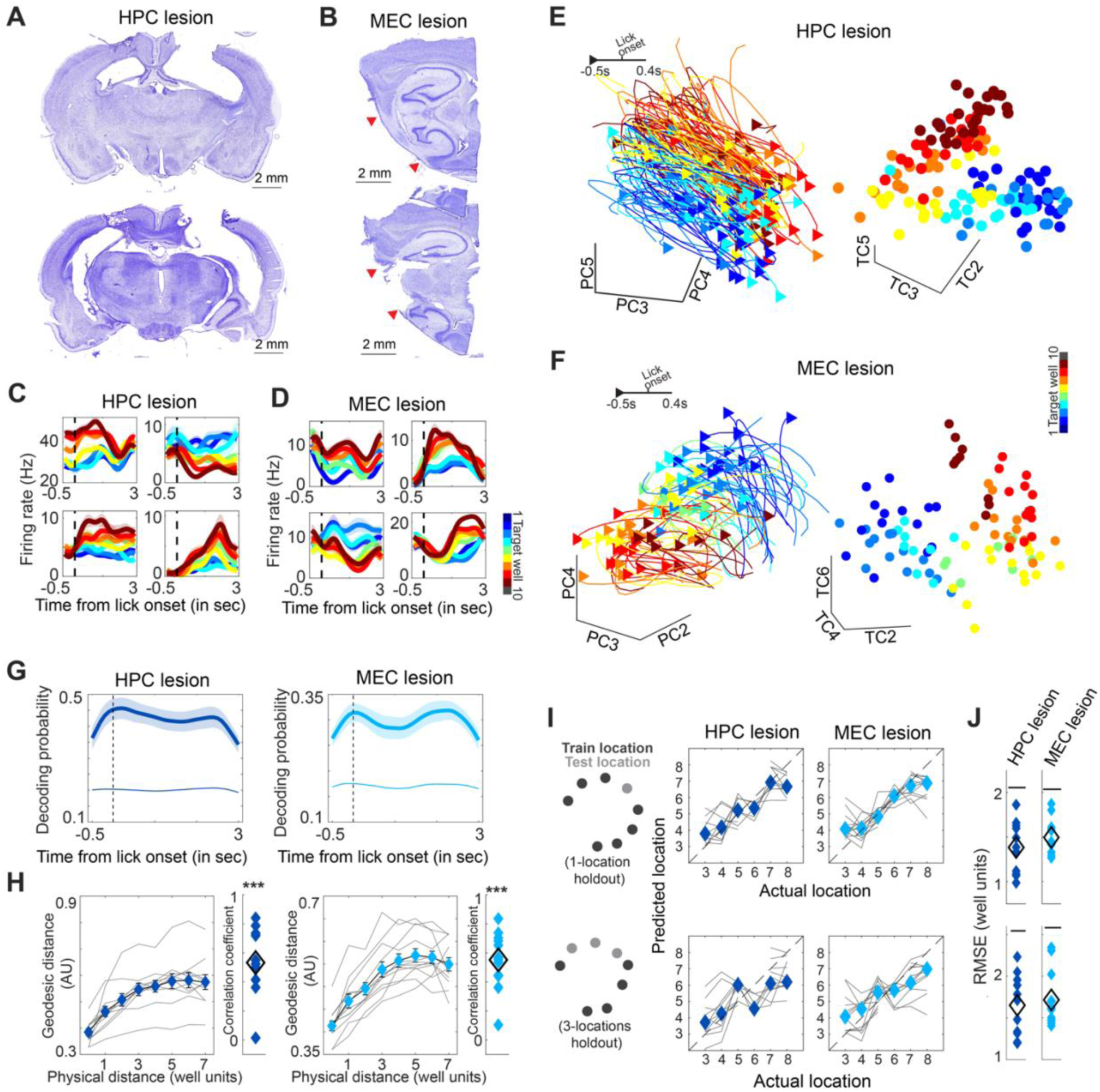
OFC spatial maps are preserved in animals with HPC or MEC lesions. (A-B) Nissl-stained sections showing HPC lesions along the dorsal-ventral axis (A) and MEC lesions (B) from a representative animal each. Red triangles indicate the region with cell body loss. **(C-D)** Target-well specific firing rates of four representative OFC neurons from animals with lesions in either HPC (C) or MEC (D). **(E-F)** OFC neural ensemble activity in a representative session from the animal with lesions in either HPC (E) or MEC (F). Left panel shows single-trial neural activity color-coded by the target well and projected on three principal components. Right panel shows the three TCA trial components of the corresponding session. Dimensions that best show the distinct representations of target wells were chosen for visualization. **(G)** Decoding probabilities of the target well identity from OFC neural ensembles of the animals with lesions in either HPC (left) or MEC (right). Shown are means (solid) ± s.e.m. (shading) across sessions. **(H)** Geodesic distances from OFC TCA trial factors in the animals with lesions in either HPC (left) or MEC (right) plotted against physical distances in well units. Each gray curve represents the median geodesic distances in a single session. Diamonds represent the means across sessions. In each panel, the inset on the right plots the correlation coefficients between median geodesic distances and physical distances for individual sessions (colored diamonds). Black open diamonds represent the means across sessions. ***p < 0.001 by Wilcoxon signed-rank test. **(I)** GPR-based prediction of target well identity from OFC trial factors from the animals with lesions in HPC (middle column) or MEC (right column). Predictions are shown for one (top) or three (bottom; either wells 3, 4, and 5 or wells 6, 7, and 8) well holdouts. Each gray curve represents the median predictions from a single session, while colored diamond depicts the median across sessions. **(J)** Root mean squared errors of well-identity predictions for the two strategies in **I**, holdout of either single (top) or three (bottom) wells. Horizontal lines represent the chance levels. Filled colored and black diamonds represent individual sessions and their means, respectively.

## DISCUSSION

In this study, we report a new form of internal spatial map in the OFC that possesses several unique features not observed in previously described maps in the HPC and MEC. Unlike the uniform mapping of locations across the environment in HPC and MEC, OFC neurons form discrete spatial representations of navigational goals in a manner that preserves their topological and distance relationships. Furthermore, when carrying out the same task in different rooms or in mazes with distinct geometries, the OFC map largely preserves the target-well representations, allowing for generalization of old maps to new contexts, in contrast to the substantial remapping observed in HPC and MEC maps under the same conditions. Below, we discuss the properties of the OFC spatial map in the broader context of the brain’s cognitive mapping system.

### Mapping of discrete positions in the OFC

Navigation often involves visiting a specific set of locations, whether it is humans moving between points of interest in an urban environment or bats in the wild traveling between particular trees^36^. In principle, such point-to-point navigation is enabled by a minimalistic spatial map that contains only the positions of interest while preserving the distance and topological relationship between them. Even in a navigation setting involving obstacle avoidance, humans^37^ and mice^38^ exhibit strategies relying on visiting key ‘subgoal’ locations to reach the destination. Consistent with this idea, neurons in the medial prefrontal cortex form a cluster of place fields selectively at relevant positions in space, either near reward-delivery sites or other navigational targets^39^. Yet, it has remained unexplored whether these neurons form a spatial map that preserves the topological arrangement between the encoded locations. Our results now confirm that OFC constructs a spatial map focused on discrete navigation targets in the maze, by forming distinct neural representations that preserve both topological and distance relationships between them (Figures 1 and 2). Under the same task conditions, we previously showed that the OFC neurons form a representation of the future navigational goal even before navigation onset, and that this representation persists throughout the entire journey^20^. These neural dynamics suggest that the primary feature of OFC ensemble activity is to segment the task into individual goal-directed journeys, each targeting a specific goal well. The initiation of navigation toward a given target triggers the corresponding spatial goal representation, which is then sustained until the animal reaches that location.

Given that all wells in our task dispense identical amounts of reward, the finding of distinct representations for individual wells may seem surprising in light of previous literature pointing to the role of OFC neurons in value encoding. In fact, some studies employing spatial tasks have suggested that OFC neural activity primarily distinguishes options based on their values^40^, resulting in overlapping representations for different positions with equivalent values^41^. In contrast, our results align with an alternative view^42^, supported by several lines of evidence, that OFC neurons encode various attributes associated with reward^43,44^, including location or accompanying action^45,34^ even when the subjective value remains identical^46,47^. In contrast to the mapping of discrete spatial landmarks in OFC, the HPC and MEC neurons track the animal’s instantaneous position by maintaining spatial continuity. As a result, these neurons in HPC and MEC maintain a largely uniform coding irrespective of whether the animal is approaching or crossing a given well (Figure 1), which is also supported by a recent study showing that place cells in HPC exhibit consistent spatial coding irrespective of whether behavior is goal-directed or random-foraging^48^.

These observations suggest that the brain possesses at least two distinct spatial mapping systems – one dedicated to maintaining a continuous spatial layout to track an animal’s instantaneous position (HPC and MEC), and another focused on behavior-relevant positions to support the planning of prospective navigation strategy (OFC)^49^. These dual mapping systems will provide substantial navigational flexibility. For example, even when the navigation target changes within the same environment, HPC and MEC can preserve a stable, continuous map to guide navigation behaviors. Conversely, when the spatial layout is altered by new barriers or obstacles, persistent target representations in OFC can still support the computation of alternative trajectories toward the same destination^50^. Flexible navigation across different spatial layouts and goals, thus likely depends on the interplay between these two mapping systems.

### Topology and distance preserved spatial mapping in OFC

The OFC has been studied primarily in the context of value-based decision making. A leading theory proposes that the OFC supports outcome inference, even under partial observability of the task structure, by constructing a ‘cognitive map’ that encodes relationships between various task components or states^51–54^. This framework accounts for classic findings, including behavioral updating following reward devaluation^55^, predicting rewards from preconditioned sensory cues^56^, and transitive inference in which relationships between paired items are inferred via their associations with a third one^57^. In line with this view, studies with functional magnetic resonance imaging (fMRI) have reported hexa-directional (grid-like) modulation in human OFC when subjects navigated an abstract social network space^58^ or a two-dimensional feature space parametrized by two independent lengths associated with an object being viewed^59^, suggesting that OFC encodes the relational structure between entities within a task space.

However, fMRI is considered to reflect pooled activity of afferent inputs within a given voxel and cannot resolve single-neuron activity, and thus, direct neuronal-level evidence for a spatial map in OFC has been lacking. Our findings provide neuronal-level evidence for a cognitive map in OFC, revealing several features not captured previously. First, the OFC forms a map of discrete behaviorally-relevant positions, rather than a uniform tiling of space, which argues against previously suggested grid-like mapping in this structure by fMRI. Second, the study confirmed that the OFC map is not limited to capturing a structure of valuable sensory cues, as previous studies have mostly focused on, but it can form a map referenced to physical space. Because all our experiments were performed in minimum light conditions, this spatial representation cannot be simply explained by the discrimination of local sensory signals associated with individual locations. Finally, the OFC map is not merely topological (graph-like) but metric, reflecting distances in the external space. This is notable because distance metrics are typically attributed to grid cells in MEC as their unique feature^60^, and even place cells in HPC are considered primarily topological^61^, rather than geometrically preserving distance and angles.

These similarities and differences between OFC, HPC, and MEC maps raise the question of their potential interdependencies. Notably, OFC spatial-map formation does not require inputs from HPC or MEC (Figure 7), implying that local computations within OFC can preserve both topology and distance of encoded locations. The absence of global remapping in OFC when recordings were performed between different rooms (Figure 4) argues against the possibility of position estimation from distal landmarks in the room. By contrast, the similar scales of the OFC maps formed for the linear and circular mazes with equidistant well-separation (Figure 5), together with the expanded map in the large maze (Figure 3), suggest that OFC neurons integrate travelled path length between locations. This hypothesis is supported by the similar increase in MEC map scale in the large maze and by prior evidence that MEC grid cells perform path integration to maintain equidistant field spacing^8,27,28^. However, the path-integration mechanism still cannot explain the OFC’s target-specific representations that persist across journeys from different start positions (i.e., different travel lengths). We thus propose that OFC neurons estimate positions as path length relative to spatial anchors in the maze. Consistent with this idea, the curved nature of the OFC spatial manifolds indicates an effective equivalence of the wells at the two extremes, suggesting that the track ends, or maze boundaries, may act as anchors from which the distance to a given well is computed based on the direction of motion.

### Neural instantiation of spatial schema

A spatial schema is an abstract representation that captures the relational structure of spatial positions shared across similar environments^1^. Behavioral studies in humans^62,63^ and animals^17,29^ suggest the use of spatial schema in navigation, and immediate-early-gene studies point to their presence in the prefrontal cortex^18,19^. Yet, direct neural evidence for a context-general spatial code has been missing. In this study, we identify a spatial schema in the OFC that generalizes across mazes of different sizes or shapes, preserving topological and distance relationships between well locations along a one-dimensional maze. OFC neurons stably encode this spatial-position arrangement with minimal remapping across environments, yielding a schematic map of well positions.

In contrast, HPC and MEC neurons undergo significant remapping across environments such that entering a new room reconfigures their population activity into a distinct neural activity subspace (Figures 4 and 6). This aligns with prior reports showing that place cells in HPC^10,9,13^, and to a lesser extent grid cells in MEC^11,64^, undergo global remapping when environments change, and that HPC neural ensemble activity distinguishes more strongly the events occurring in different rooms, compared to those within the same room^65^. While remapping is useful for discriminating between environments, its sensitivity to environmental change poses a challenge for generalizing task knowledge across rooms. In fact, multiple lines of behavioral evidence, including our present results, show that the animal can generalize spatial tasks across rooms - once animals are trained in one room, they can perform the same task immediately in a new room. Moreover, while maps in HPC and MEC capture fine-grained spatial details, schematic maps are supposed to abstract common spatial structures across environments. For example, a patient with HPC damage could still recall a schematic layout of a familiar environment despite losing specific spatial details^66^. These results together suggest that spatial schemas are unlikely to be supported by HPC or MEC^1^.

To probe spatial schema by neural recordings, it is essential to dissociate the neural coding of spatial position from the representation of task progress. In fact, the navigation phase is a key behavioral variable represented in the OFC. The OFC neural population activity evolves with a similar dynamics from the journey onset to goal arrival, whereas the target-location information resides in a subspace orthogonal to these dynamics^20^. Conventional T-maze and other fixed-goal navigation tasks confound this navigation-phase representation with the position coding, hindering this dissociation. By requiring animals to reach the same wells from multiple start positions and approach directions, our task design can dissociate position from navigation phase, enabling identification of allocentric spatial coding in a linear-maze task. Furthermore, the comparative analysis of spatial maps across OFC, HPC, and MEC carried out in multiple environments contrasts the key differences between these three mapping systems, allowing us to conclude that the spatial schema across environments is a unique feature of the OFC map. Although previous studies have suggested that HPC plays a key role in supporting the formation of spatial schema in the prefrontal cortex^2,17^, our results indicate that the OFC spatial schema can emerge independently of inputs from HPC or MEC. We note, however, that our one-dimensional task imposes modest spatial demands, and in more complex environments, the fine-grained spatial layouts supplied by HPC or MEC may be necessary for OFC schema formation^2,17^.

The spatial schema we observed in OFC aligns with a broader class of generalizable coding schemes observed in this brain region. In tasks requiring specific actions at individual locations according to a sequence of odor cues, OFC neurons encode the ordinal positions even when sets of odors were exchanged^16^. In a value-based decision task, OFC neurons encode the value of offered and chosen options independent of option identities^67^, and in a perceptual decision task, the confidence representation in OFC is invariant to sensory modalities^68^. These observations point to a role for the OFC in extracting task-relevant structure while compressing irrelevant details.

Flexible behavior and learning require not only generalization but also context-specific specialization. We propose a division of labor in which OFC supplies an abstract context-general schema, while HPC and MEC maintain detailed context-specific maps for solving the ongoing problem. This idea is supported by a computational model in which learning is accelerated in a recurrent neural network with both problem-general and specific components^69^. Interactions between these two cognitive maps can retune population dynamics with minimal network changes, by reusing a learned task-relevant neural manifold within a new task context, likely underlying the brain’s hallmark capacity for rapid and efficient behavioral adaptation and learning.

## METHODS

### Subjects

All experiments were approved by the local authorities (RP Darmstadt, protocols F126/1009 and F126/1026) in concordance with the European Convention for the Protection of Vertebrate Animals used for Experimental and Other Scientific Purposes. Sixteen male Long-Evans rats weighing 400 to 550 g (aged 3–6 months) at the start of the experiment were housed individually in Plexiglass cages (45×35×40 cm; Tecniplast GR1800) and maintained on a 12h-light/12h-dark cycle, with behavioral experiments performed during the dark phase. All animals were water-restricted with unlimited access to food and kept at 90% of their free-feeding body weight throughout the experiment. Eleven rats had tetrodes implanted only in the OFC, out of which two received bilateral lesions in the dorsal and ventral HPC, and two received bilateral lesions in the MEC. Four of the non-lesioned rats with tetrodes in the OFC were used in a previous publication^20^. Two rats had tetrodes implanted both in the OFC and the dorsal HPC. One rat had tetrodes implanted only in the dorsal HPC. Two rats had tetrodes implanted in the dorsal MEC. No statistical method was used to predetermine the sample size.

### Surgery, lesion injection, and drive implantation

Anesthesia was induced by isoflurane (5% induction concentration, 0.5–2% maintenance adjusted according to physiological monitoring). For analgesia, Buprenovet (Buprenorphine, 0.06 mg/mL; WdT) was administered by subcutaneous injection, followed by local intracutaneous application of either Bupivacain (Bupivacain hydrochloride, 0.5 mg/mL; Jenapharm) or Ropivacain (Ropivacain hydrochloride, 2 mg/mL; Fresenius Kabi) into the scalp. Rats were subsequently placed in a Kopf stereotaxic frame, and an incision was made in the scalp to expose the skull. After horizontal alignment, several holes were drilled into the skull to place anchor screws, and craniotomies were made for microdrive implantation. The microdrive was fixed to the anchor screws with dental cement, while two screws above the cerebellum were connected to the electrode’s ground. All animals received analgesics (Metacam, 2 mg/mL Meloxicam; Boehringer Ingelheim) and antibiotics (Baytril, 25 mg/mL Enrofloxacin; Bayer) for at least 5 days after the surgery.

For tetrode recordings, rats were implanted with a microdrive that contained individually adjustable tetrodes made from 17 µm polyimide-coated platinum-iridium wire (90–10%; California Fine Wire; plated with gold to impedances below 150 kΩ at 1 kHz). The tetrode bundle consisted of 30-gauge stainless steel cannula, soldered together in circular or rectangular shapes. Four rats with drives implanted in the OFC (Rat 110, Rat 175, Rat 182, and Rat 284) have been described previously^20^. Rats 499, 541, 475, 519, 531, and 647 were implanted with two circular bundles, each with 14 tetrodes, targeting the OFC in both hemispheres. Rat 419 was implanted with a drive targeting the OFC bilaterally, and comprising two circular bundles with 16 tetrodes each. Rats 497 and 533 were implanted with drives that had circular bundles targeting both the OFC and the dorsal HPC bilaterally. In these animals, the OFC bundles comprised 16 tetrodes each. In rat 497, bundles with nine and four tetrodes were implanted over the left and right HPC, respectively. In rat 533, bundles with 11 tetrodes each were implanted over the dorsal HPC. Rat 666 was implanted with a drive targeting the dorsal HPC in both hemispheres using two bundles with 14 tetrodes each. Rats 571 and 606 were implanted with two circular bundles with 14 tetrodes each, targeting the MEC bilaterally. All bundles targeting OFC were centered at 3.75 mm anterior-posterior (AP) relative to bregma, and 2.5 mm medial-lateral (ML) orthogonal to the AP axis. Bundles targeting the dorsal HPC were centered at 3.8 mm AP (posterior to bregma) and 3 mm ML. Bundles targeting the MEC were centered at 0.2 mm anterior to the transverse sinus, and 4.5 mm ML and implanted at 20° relative to the vertical. Experiments began at least 1 week after the surgery to allow the animals to recover.

For lesion surgeries, we used Ibotenic acid (Hello Bio), dissolved in phosphate-buffered saline at a concentration of 10 mg/ml at pH 7.5. For lesioning the entire dorsal and ventral HPC bilaterally, we injected Ibotenic acid at 13 sites per hemisphere, based on previously described set of coordinates^70^ (AP, ML, and DV in mm:-2.4, 1.0, 3.4;-3.0, 1.4, 2.6*;-3.0, 1.4, 3.4*;-3.0, 3.0, 3.0;-4.0, 2.6, 2.3*;-4.0, 2.6, 3.3*;-4.0, 3.7, 3.0;-4.9, 3.9, 3.5*;-4.9, 3.9, 7.0*;-5.7, 4.1, 3.8;-5.7, 5.1, 4.0;-5.7, 5.1, 4.9;-5.7, 5.1, 5.8). A volume of 100 nl was injected at each site, except the locations marked with *, where 50 nl was injected. For bilateral MEC lesions, we injected 100 nl of Ibotenic acid at three sites per hemisphere at an angle of 20° relative to the vertical (AP relative to the transverse sinus, ML, and DV in mm: 0.2, 4.6, 4.5; 0.2, 4.3, 2.5; 0.2, 4.3, 3.5). The drug was injected with an infusion rate of 50 nL/min using a 1 μl Hamilton syringe (7000 series). After the injection was completed, the needle was left in place for 5 min. Behavioral training was started at least two weeks after the lesion surgery, and tetrode drives were implanted at least 4 weeks after the lesion surgery. Neural recordings started at least 1 week after the tetrode implantation.

### Behavioral methods

Rats were trained in a two-meter-long linear maze with 10 reward wells distributed at an equal distance (20 cm) between each other. The training procedure proceeded in three phases as described in detail previously^20^. Briefly, in the first phase, 100 μl of liquid reward (0.3% saccharin) was delivered manually at two specific reward wells in an alternating manner. Once rats started drinking the water reward from the wells, they entered the second training phase, where a liquid reward was delivered only after they licked the correct well. The delay between the initial lick detection and the reward delivery was gradually increased to 1-2 seconds. Finally, in the third phase, the reward-well transition rule was introduced, which required animals to update their targets to new pairs of rewarded wells after several successful trials. Transitions were signaled by white LEDs positioned underneath all ten reward wells as well as the delivery of the liquid rewards in the two new locations. The behavior analyses (Figure S9D) started from the first day of phase 3. Reward well pairs were chosen such that a given well is approached from different starting locations and directions. In addition, adjacent wells were never chosen as rewarding well pairs. The animal’s position and head direction were monitored with two-colored LEDs on the head stage at a sampling rate of 25 Hz. All the recordings were performed under a minimum-light condition (no light source in the recording room, only with weak ambient light coming from the adjacent room with computer monitors).

In most neural recording sessions in the linear maze, we used all wells from well 2 to well 9. Wells 1 and 10 were used in only one session and were generally avoided in order to prevent the animals from using the maze ends as a cue to visit these wells. In 14 sessions, we withheld reward delivery at one or more of the middle wells in order to check how non-rewarding wells affected the neural representations of adjacent wells (Figure S5A).

Paired sessions were performed only when the animals were sufficiently trained in the alternation task in the linear maze located in one room. Three additional mazes were used for the paired sessions. To study the scaling of the spatial maps (Figure 3), a 4-meter-long semicircular maze was used, with ten equally distributed reward wells, spaced 40 cm apart. To study the impact of room changes on spatial representations, we used an identical 2-meter-long linear maze located in a different room (Figure 4). The new room (room 2) was smaller in size than room 1, along with different distal cues. For example, one of the walls was replaced by a curtain in room 2, and the two rooms had different wall installations. To align the well identities of the two mazes, we used two sets of cues - we positioned a wooden board above well 2 as a local cue in both mazes - thus only wells from 3 to 9 were used in the paired experiments involving the two rooms - and as a distal cue, the experimenter and the computer were positioned on the same side relative to the mazes in both rooms. The ceiling lights were turned off once the animal was placed in the mazes, and the recordings were performed in minimal light conditions. Finally, to study the effect of maze geometry on spatial mapping, we used a circular maze with a 2-meter circumference (Figures 5 and 6). The well separation in the circular maze was 20 cm, identical to that in the linear maze. In addition, a barrier was placed between wells 1 and 10 to make the linear and the circular mazes topologically identical. The equivalence of the well-identity assignment across the two tracks was based on the relative positions of the wells. Both mazes were positioned such that, from the perspective of the experimenter, well 9 is to the left of well 2. For all paired-session experiments, animals were habituated in a new maze for 1-2 sessions before starting neural recordings. Importantly, animals did not require extensive retraining and could readily perform the alternation task in the new maze (Figures S6G and S8J). The order of the two mazes was selected in a pseudo-random manner.

### Histological procedures

Once the experiments were completed, the animals were deeply anaesthetized by sodium pentobarbital and perfused intracardially with saline, followed by 10% neutral buffered formalin solution. The brains were extracted and fixed in formalin for at least 72 hr at 4 °C in temperature.

Frozen coronal sections were cut (30 µm) and stained using cresyl violet and mounted on glass slides.

### Spike sorting

All data processes and analyses were performed with MATLAB (MathWorks). Neural signals were acquired and amplified using two or four 64-channel RHD2164 headstages (Intan Technologies), combined with an OpenEphys acquisition system at a sampling rate of 15 kHz. The signals were band-pass filtered at 0.6-6 kHz, and spikes were detected and assigned to separate clusters using Kilosort^71^ (https://github.com/cortex-lab/KiloSort) under the parameter settings of the spike threshold at-4 and the number of filters at 2 times the total channel number. Each tetrode was independently grouped with ‘kcoords’ parameters, and the noise parameter determining the fraction of noise templates spanning across all channel groups was set to 0.01. The obtained clusters were checked and adjusted manually based on autocorrelograms and waveform characteristics in principal component space, obtaining well-isolated single units by discarding multi-unit activity or noise. Neurons with firing rates less than 0.5 Hz were excluded. The firing-rate estimation was performed by convolving spike times with a Gaussian kernel with a bandwidth of 250 ms, except during spatial firing rate computation (see ‘Spatial firing rates and spatial information’ below).

### Spatial firing rates and spatial information

To compute the spatial firing rates, we binned the linear track into 2.5 cm bins and calculated the firing rate in each bin by only considering spikes during periods of motion (>20 cm/s). The raw firing rate was further convolved with a 25 cm wide (10 bins) Gaussian filter. The preferred running direction for a given neuron was considered to be the direction with the higher peak firing rate. Spatial information (Figures S2A and S8C) was obtained from the rate maps using the following formula (Skaggs et al., 1996),

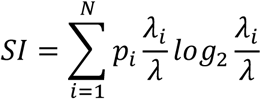

where 𝑁 is the total number of spatial bins, 𝑝_𝑖_ is the probability of occupying the i^th^ bin, 𝜆_𝑖_ is the firing rate in the i^th^ bin, and 𝜆 is the overall average firing rate of the neuron.

### Cell selection

No cell selection was involved for the analyses of distinct spatial coding schemes between target and non-target wells during goal-directed journeys (Figures 1 and S2). For the analysis of spatial map structures during approach and licking (Figures 2-7 and S3-S9), we selected neurons that exhibit target well-specific firing during-0.5 s to 3 s relative to the onset of licking, based on the following criterion: Firing rates of individual neurons were segmented into 100-ms bins. As OFC neurons exhibit highly dynamic firing rates during approach and licking (Figure S3), we factored out the temporal firing dynamics shared across all trials to extract trial-wise contributions of a given neuron. For each neuron, we constructed the firing rate matrix **F** with dimensions K × T, where K is the number of trials and T is the number of time bins (36 bins). Next, we mean-centered the matrix and performed singular value decomposition (SVD, Figures S3A-D), which factorizes the matrix **F** as follows,

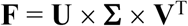

**U** and **V** are orthonormal matrices with dimensions K × r and T × r, respectively. **Σ** is a diagonal matrix of dimension r × r, where r is the rank of matrix **F**. We observed that reconstructing the matrix **F** using only the first columns of the **U** and **V** explained a median of 44.9%, 57.9% and 51.6% of the variance in firing rates of OFC, HPC, and MEC neurons, respectively (Figure S3D). Hence, we considered the first column of **U** (a K × 1 vector) as trial-wise contributions of an individual neuron within a session, while the first column of **V** represents major temporal firing dynamics over T time bins. Subsequently, one-way ANOVA was performed on trial-wise contributions with the identity of target well as an independent factor. Cells with p < 0.05 in the ANOVA test were considered as target-well selective.

In paired sessions, the firing rate matrix F was constructed by concatenating the activity over the two sessions, although the ANOVA was performed separately for each session. Cells that were significantly well-selective in at least one of the two sessions were considered for further analysis.

### Tuning distance and ΔTS computation

We first computed the relative coding strength across different target wells for a given neuron by regressing the trial-wise contributions (described in the previous section) using the target-well identity as a categorical predictor, shown in the following expression,

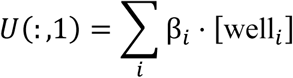

Where β_i_ is the coding strength for the i^th^ target well, and [well_i_] is a one hot coding vector. The regression model was fitted using the MATLAB function ‘fitlm’. No intercept term was included as the firing rate matrix F was mean-centered prior to SVD. Hence, the regression coefficients are essentially the mean of trial-wise contributions for a given target well. The β vector, comprised of the β_i_s, is defined as the coding strength vector for the given neuron (Figure S3G). Importantly, in paired sessions, a separate β is obtained for each session. Subsequently, tuning distance was computed as the cosine distance between β**^session1^** and β**^session2^**, defined as 1-cos(θ) where θ is the angle between β^session1^ and β^session2^. ΔTS was computed as (β^session2^ - β^session1^)/ (β^session2^ + β^session1^).

### Population vector analysis

A population vector was constructed for each trial, comprising the trial-wise contribution from all well-selective neurons. The representational distance between two trials was computed as the cosine distance between the corresponding population vectors. To quantify the difference between spatial maps in paired sessions, we computed the median cosine distance between all trial pairs belonging to well ‘i’ and well ‘j’. This well-specific representational distance was calculated for well-pairs both within (𝑑𝑖,𝑗^self^) and across sessions (𝑑𝑖,𝑗^other^). Next, we computed the difference between d^other^ and d^self^, averaged over all well pairs and normalized to the sum as follows,

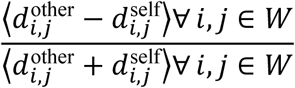

Where 〈.〉 denotes the mean, and W is the set of reward wells used in the paired session.

### Dimensionality reduction using PCA and TCA

Before implementing PCA to reduce dimensions of neural ensemble activity, we implemented a soft-normalization technique described by Churchland et al.^72^, to adjust the range of firing rates. To perform PCA on neural activity during-0.5 s to 3 s relative to lick onset, we constructed a data matrix with K*T rows and N columns, where K, T, and N denote the numbers of trials, time bins (36 times for a 3.5 seconds duration), and well-selective neurons, respectively. For paired sessions, trials from the two sessions were concatenated to generate a set of PC axes common to both sessions. PCA was performed on the data matrix using the MATLAB Toolbox for Dimensionality Reduction (https://lvdmaaten.github.io/drtoolbox/).

To perform TCA, we used denoised neural data, defined as the neural activity along a set of PC dimensions that explain 85% of the data variance. The data tensor had a dimension of K × T × P, where P is the number of PC dimensions used. TCA was performed using the higher-order singular value decomposition algorithm (HOSVD^73^) implemented using the MATLAB toolbox ‘tensortoolbox v3.2’ (https://www.tensortoolbox.org/). TCA shrinks the tensor along each of the data dimensions, and in this paper, we focused on the trial factor matrix, whose rows reflect a unique population-level contribution to a given trial. The trial factor matrix is an orthonormal matrix with dimensions K × r, where r (trial components) can be defined by the user. Importantly, the columns of the trial factor matrix are ordered according to the variance (2-Norm) described in the original tensor, thereby giving greater importance to the first few trial components. In Figure 2G, we show that target-well decoding performance improved rapidly as a greater number of trial components were considered, and we found that 8, 6, and 6 trial components from the OFC, HPC, and MEC, respectively, achieved 85% of the saturating target well decoding levels. Hence, the target-well identity is a major contributor to trial-wise variances of neural population activity in all three brain regions. In paired sessions involving the large track, 11, 6, and 8 trial components from the OFC, HPC, and MEC, respectively (as used in Figure 3F), explained on average 82.77%, 85.43%, and 87.88 % of the saturating well decoding levels when combined across both tracks. In paired sessions involving the circular track, 10, 5, and 6 trial components from the OFC, HPC, and MEC, respectively (as used in Figure 5D) explained 83.51%, 81.28%, and 80.62 % of the saturating well decoding levels when combined across both tracks. While the main figures present the results of TCA-based analyses using specific numbers for trial components, Figures S4D-E, S5E-G, S6H, and S8H-I show the results by using a range of trial component numbers.

### Geodesic distance computation

For a given number of trial components, we first constructed a nearest neighbor (NN) graph of the trial factors (nodes in the graph) using a modified version of the original Isomap algorithm^74^. The graphs are usually constructed by connecting each node to a specified number of nearest neighbors (NN = 6 for the analyses in the main figures), and the geodesic distance is computed as the shortest distance between two nodes when traversing along the graph using Dijkstra’s algorithm. However, if some of the nodes are tightly clustered, for example, when trials targeting the same wells have very similar representations, the nearest neighbor graph may end up containing disconnected components with no way of reaching a node in one component from a node in the other. Hence, in such scenarios, we slightly modified the graph construction algorithm by identifying a disconnected pair of components, and manually connecting the pair of nodes, one from each component, that are nearest to each other. This modification was performed until there were no disconnected components, thereby allowing geodesic distance measurement between any two nodes. Although such modifications can be circumvented by using large NNs, we show in Fig. S4F that such strategies would ignore the fine-scale nonlinearities within the manifold, thereby obfuscating the near-linear relationships between representational geodesic distance and physical distance. Pearson’s correlation coefficients were computed between the median geodesic distances of all trial pairs targeting a given pair of reward wells and their physical distances in well units (distances between trials with the same target wells were excluded). One well unit corresponds to the distance between two adjacent wells.

To verify topological maps at different time points during approach and licking, we performed the geodesic distance analysis at-0.5s, 0s, 1s, 2s, and 3s relative to lick onset (Figure S3I-S3J). PCA was performed on neural data across trials for a given time point relative to lick onset, and geodesic distance was computed on 8, 6, and 6 PC dimensions from OFC, HPC, and MEC neural activity, respectively, in order to match the analysis performed on trial components obtained post-TCA (Figure 2J).

### Gaussian process regression

GPR enables regression over a curved manifold as long as the dependent variable varies smoothly with the predictors. Let (**y**, **x**) be a set of observations, where **y** is the animal’s location, and **x** is the corresponding neural population activity. **y** can be considered as a function **f** of **x**, and is modeled as a Gaussian process with zero mean and a covariance matrix whose entries are defined by a squared exponential kernel as follows,

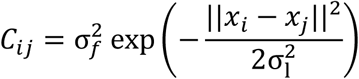

where σ_𝑓_and σ_𝑙_ are hyperparameters fit by the model and **x_i_** and **x_j_** are any two neural activity vectors. The squared exponential kernel ensures that **y** varies smoothly over **x** and that nearby **x**’s correspond to similar y values. Once the Gaussian process model is built, the regression problem can be framed as predicting **f(x*)** given a query data point **x*,** which is not a part of the original data matrix **X**. **f(x*)** is estimated as the conditional distribution of the multivariate Gaussians evaluated at **x\***. Finally, the expectation of **f(x*)** is considered to be the predicted location of the animal for the given neural activity **x\***. The algorithm is implemented using MATLAB’s ‘fitrgp’ function.

In this paper, GPR was used for two sets of analysis. In Figure 1C-1D, we used GPR to decode the animal’s location, specifically during the periods of goal-directed journey when the animal’s speed was above 20 cm/s. For this decoding analysis, we used denoised neural data obtained by performing PCA on the neural activity over the entire session, and then choosing the PC dimensions that explain 85% of the variance. Next, to account for the difference in firing rates depending on running direction, we divided the entire data set into two parts. We trained a simple LDA-based decoder (a binary classifier) on one part of the data to decode the animal’s direction of movement. This decoder was subsequently used on the other part to predict the movement direction at each 100-ms time bin. For each predicted direction, we separately performed a 10-fold cross-validated GPR-based location prediction. This method ensured that the direction-based GPR modeling was in accordance with the degree of direction separability of the neural data.

The second set of GPR-based analysis was designed to predict the target-well identity from TCA trial components. Importantly, the analysis considered well identities as a continuous, rather than categorical, variable, which enables a regression-based analysis (See the section on ‘Decoding analysis’ for analyses where well identities are treated as categorical variables). To perform a GPR analysis with a single held-out well, we excluded all trial components associated with that well from the training set. For the 3-well hold-out GPR, we excluded trial components associated with either wells 3, 4, and 5 or wells 6, 7, and 8 for training the GPR model. The held-out trial components were used for prediction. Because GPR is most effective for interpolation, rather than extrapolation, we did not hold out the end wells (2 and 9) for prediction. For the 3-well hold-out analysis, we considered only those sessions in which both well 2 and well 9 were used. To compute the chance levels for well-identity prediction, we shuffled the class labels (well identities) 100 times during the GPR training. For each shuffle, we computed the RMSE of the predicted well identities based on the held-out trial factors. The 5^th^ percentile of the resulting RMSE distribution was taken as the chance level.

### Decoding analysis

#### Decoding well identities during crossing and licking

To compare spatial coding during approach and licking a target well versus crossing a non-target well, we trained two sets of decoders – a cross-well decoder, trained on the identities of crossed wells, and a target-well decoder, trained on the identities of target wells during approach and licking. The neural activity used for decoding was denoised by choosing the PC dimensions that explain 85% of variance over the entire session. To train the cross decoder, we used neural activity from three consecutive 100-ms bins preceding the animal’s crossing of a well during a goal-directed journey. Only well-crossing events at speeds greater than 20 cm/s were included. The cross-decoder was tested using a leave-two-out cross-validation approach, where, for the prediction of the well identity in a given journey, all well-crossing events from the ongoing and immediately preceding journeys were excluded from the training dataset. The same cross-validation technique was implemented when using the cross-decoder to predict target-well identity during approach and licking. For the target-well decoder, we used neural activity from 0.5 s before to 3 s after the lick onset of a target well. Only rewarded trials were included to train the target-well decoder, and decoding was performed using a leave-two-out cross-validation approach.

To account for differences in firing patterns based on running direction^24^, we used a Gaussian mixture model discriminant analysis (GMMDA) decoder^25^. GMMDA models the neural population activity corresponding to a particular class label as a mixture of *n* multivariate Gaussian subclasses, such that the probability density function of class *k* is defined as,

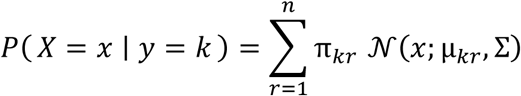

Where **x** is a neural activity vector, 𝜋_𝑘𝑟_ and 𝝁_𝒌𝑟_ represent the mixing probability and the mean of the r^th^ subclass of class *k*, respectively, and 𝚺 is the common covariance matrix. The parameters, 𝜋_𝑘𝑟_, 𝝁_𝒌𝑟_, and 𝚺, were estimated using the expectation-maximization algorithm. We used 2 subclasses (n = 2) to model the possible separation of neural activity depending on the two running directions. Although it is possible to make the model more flexible with different subclass numbers for individual classes or by using distinct covariance matrices for each class and subclass, our approach with the same subclass number and shared covariance matrix across classes reduces the number of parameters required to fit the model. Once the model parameters were estimated, the decoding was performed by estimating the posterior 𝑃( 𝑦 = 𝑘 ∣ 𝑿 = 𝒙), by applying Bayes’ rule to the above expression. For well-identity estimation, we first implemented the multiclass GMMDA in a pairwise configuration, training binary classifiers for individual well pairs. The posterior of well identity was then derived from the pairwise posteriors using an algorithm described by Hastie and Tibshirani^75^. GMMDA-based decoding resulted in significantly higher decoding probabilities compared to LDA (Figure S2E-F). GMMDA was implemented using custom-written scripts and MATLAB’s ‘predict’ function.

#### Subspace-based decoding

The subspace-based decoding analysis examines how target-well information in a spatial map from one context is embedded within the subspace – a set of axes that captures the majority of neural variance, and its corresponding nullspace of a spatial map from another context. By projecting neural activity from one context to the subspace and nullspace defined in another context, we estimated the extent to which target-well information is shared between contexts. For a fair comparison between the two projections, we ensured equal dimensionality of the subspace and the nullspace by following these steps. First, for a given paired session, we constructed two data matrices, one for each session, comprising neural activity during approach and licking as described in the previous section on ‘Dimensionality reduction using PCA and TCA’. Next, we performed PCA on one of the sessions and defined the nullspace as the set of final PC dimensions that explain the last 10% of the data variance. The subspace was subsequently defined as an equal number of leading PC dimensions. If the neural activity of a given session was highly low-dimensional, such that more than half of the total PC dimensions are required to explain the last 10% of the data variance, we instead divided the total number of PC dimensions into two equal halves, with the subspace and the nullspace occupying the top and bottom halves, respectively. Overall, the subspace dimensions explained a median of 87.21% of the data variance while comprising a median of the top 47.89 % of PC dimensions across all brain regions. As a positive control, we found that the projecting neural activity on its own nullspace yielded very poor decoding performance of target-well identity, in contrast to the high decoding performance observed when projecting on its own subspace (see ‘Same sub’ versus ‘Same null’ in Figures 4G and 6G). Once the subspace and the nullspace for a given session were defined, the neural activity from the paired session was projected on these spaces, followed by target-well decoding on the projections. Decoding was performed using a multiclass LDA-based classifier with a leave-one-out cross-validation strategy, implemented in MATLAB using the ‘fitcecoc’ and ‘predict’ functions. Subspace similarity index for a given paired session was defined as the difference in target-well decoding probability in the ‘Other’s subspace’ and the ‘Other’s nullspace’, normalized by the corresponding difference between the ‘Same subspace’ and the ‘Same nullspace’, and then averaged across the two sessions and over the decoding duration.

#### Target-well decoding using TCA trial components

Target-well identity was predicted from the trial components derived from TCA using an LDA-based decoder with 10-fold cross-validation. To directly compare decoding performance across varying numbers of trial components, the same cross-validation set was used for a given session.

#### Chance level calculation

Chance levels calculation was performed as described in Basu et al.^20^. Briefly, we observed that the animal’s running direction was the largest confounding factor when decoding target-well identities. Hence, we divided the training dataset into two groups based on the approach direction and shuffled the class labels within each group. Shuffling was performed 100 times, and for each shuffle, we computed the target-well decoding probability averaged over all trials. The chance level for a given session was defined as the 95^th^ percentile of this shuffled distribution. This shuffling strategy preserved the direction-based separation in neural activity, and hence, decoding performance above the chance level indicates the well-identity information beyond that explained by movement direction. To obtain a chance level across sessions, we randomly chose one of the 100 shuffled average decoding probabilities from each session and computed their mean. Next, this process was repeated 1000 times, and the 95^th^ percentile of the distribution of means was considered the aggregate chance level.

### Visualization

#### LDA-based visualization of cross-and target-well subspaces

For the visualization in Figure 1E, we focused on 300 ms (three consecutive 100-ms bins) of neural activity preceding the crossing of an intermediate well during a goal-directed journey and 300 ms of neural activity preceding the onset of licking a target well during rewarded trials. The neural activity comprised the top PC components that explained 70% of the data variance over the entire session. For computing the cross-and target-well subspaces, we performed the supervised dimensionality reduction technique LDA (implemented using the ‘lda’ function in the MATLAB Toolbox for Dimensionality Reduction) while using the corresponding well identities as class labels. We orthogonalized the first two LDA dimensions (using MATLAB’s ‘orth’ function) and projected the neural activity on these dimensions for visualization. Similarly, to visualize the target-well activity on the cross-well subspace, we projected the neural activity during approach on the orthogonalized LDA dimensions obtained during crossing.

#### UMAP-based visualization

For visualizing the neural manifold during motion (Figure 1C), we first segmented the neural activity time series into 100-ms bins and denoised it by taking the PC dimensions that explain 85% of the data variance over the entire session. Next, we performed UMAP on the binned and denoised neural activity during the entire session, and used only the time bins during goal-directed journeys along a given motion direction, colored according to the animal’s location, for visualization. UMAP was implemented using MATLAB’s ‘run_umap’ function with 50 ‘n_neighbors’ and 3 ‘n_components’.

## ACKNOWLEDGEMENTS

We thank N.Vogt, S. Zeissler, E. Northrup, and G. Wexel for animal care; F. Bayer and A. Umminger for building the behavioral mazes; and all members of the Ito and Basu laboratories for discussions. This work was supported by the Max Planck Society (H.T.I), The Edmond and Lily Safra Center for Brain Sciences (R.B.), The European Research Council Grants (‘ChoiceSpace’ Grant Agreement no. 101117542 to R. B.; ‘NavigationCircuits’ Grant Agreement no. 714642 and ‘MentalTravel’ Grant Agreement no. 101087404 to H.T.I.), and the Abisch-Frenkel Foundation (R.B).

## AUTHOR CONTRIBUTIONS

R.B. and H.T.I. conceptualized the study and designed experiments. R.B., M.M., I.B., and C.U. performed all experiments. R.B. performed all data analysis. R.B., and H.T.I. wrote the manuscript after discussion among all authors. R.B. and H.T.I. co-supervised and coordinated the project.

## DATA AVAILABILITY

The datasets used in this study will be deposited in the Dryad database.

## CODE AVAILABILITY

MATLAB codes for PCA, LDA, and Isomap are part of the ‘dimensionality reduction toolbox’ written by Laurens van der Maaten (https://lvdmaaten.github.io/drtoolbox/). TCA was implemented using the MATLAB toolbox ‘tensortoolbox v3.2’ (https://www.tensortoolbox.org/). Other codes are publicly available at https://doi.org/10.5281/zenodo.17102288.

