## Supplementary Figures 1-9 for "The orbitofrontal cortex forms a context-generalized spatial schema that preserves topology and distance"

**The PDF file includes:**

Supplementary figures S1-S9

**Figure S1**

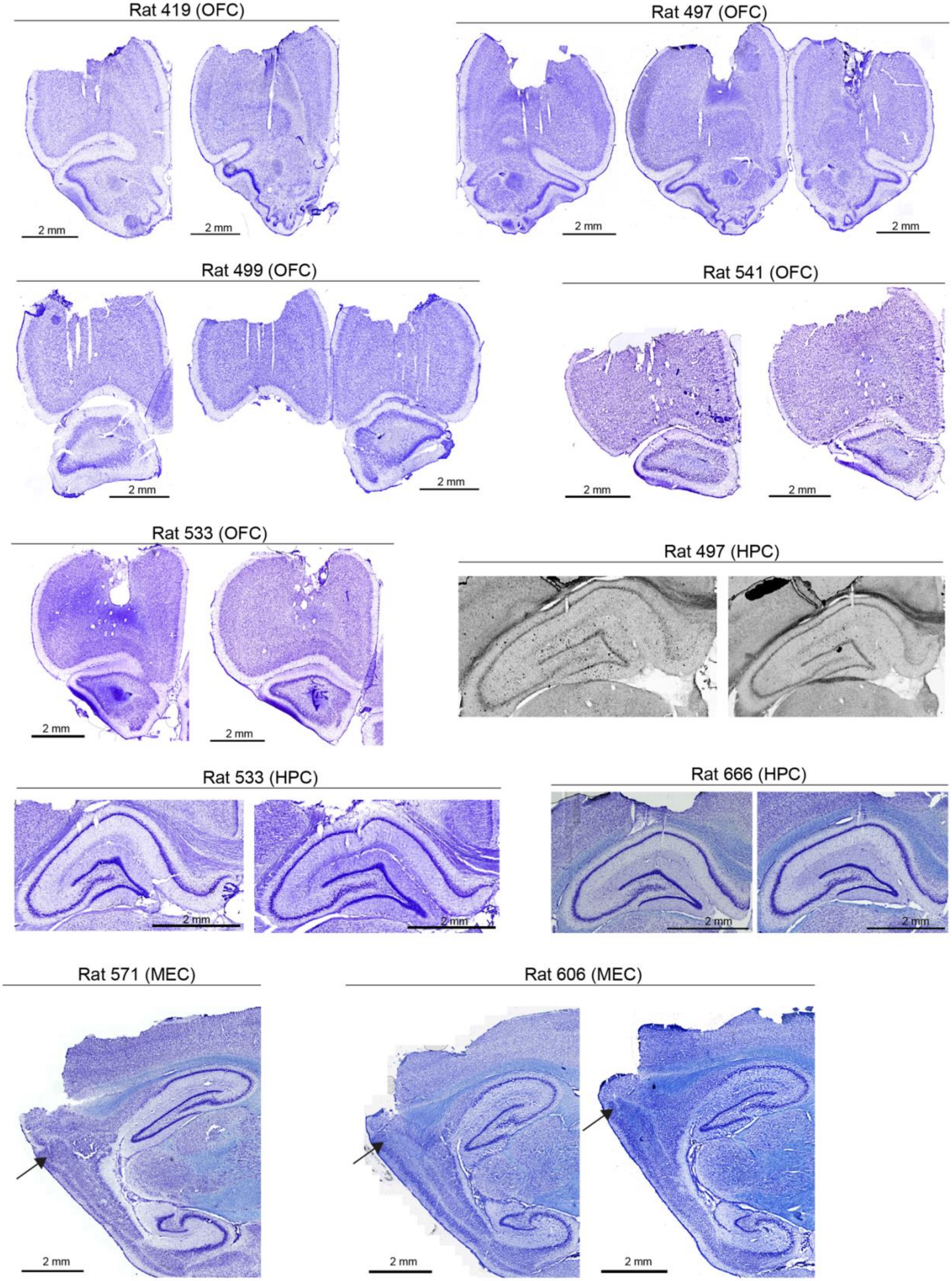
Rat 419 (OFC)
2 mm
2 mm
Rat 497 (OFC)
2 mm
2 mm
Rat 499 (OFC)
2 mm
2 mm
Rat 541 (OFC)
2 mm
2 mm
Rat 533 (OFC)
2 mm
2 mm
Rat 497 (HPC)
Rat 533 (HPC)
2 mm
2 mm
Rat 666 (HPC)
2 mm
2 mm
Rat 571 (MEC)
2 mm
Rat 606 (MEC)
2 mm
2 mm

**Figure S1: Histological sections (related to Figures 1-6):** Nissl-stained brain sections highlighting tetrodes targeting in the OFC, hippocampal CA1, and dorsal MEC. Arrows point to the tetrode locations.

### Figure S2

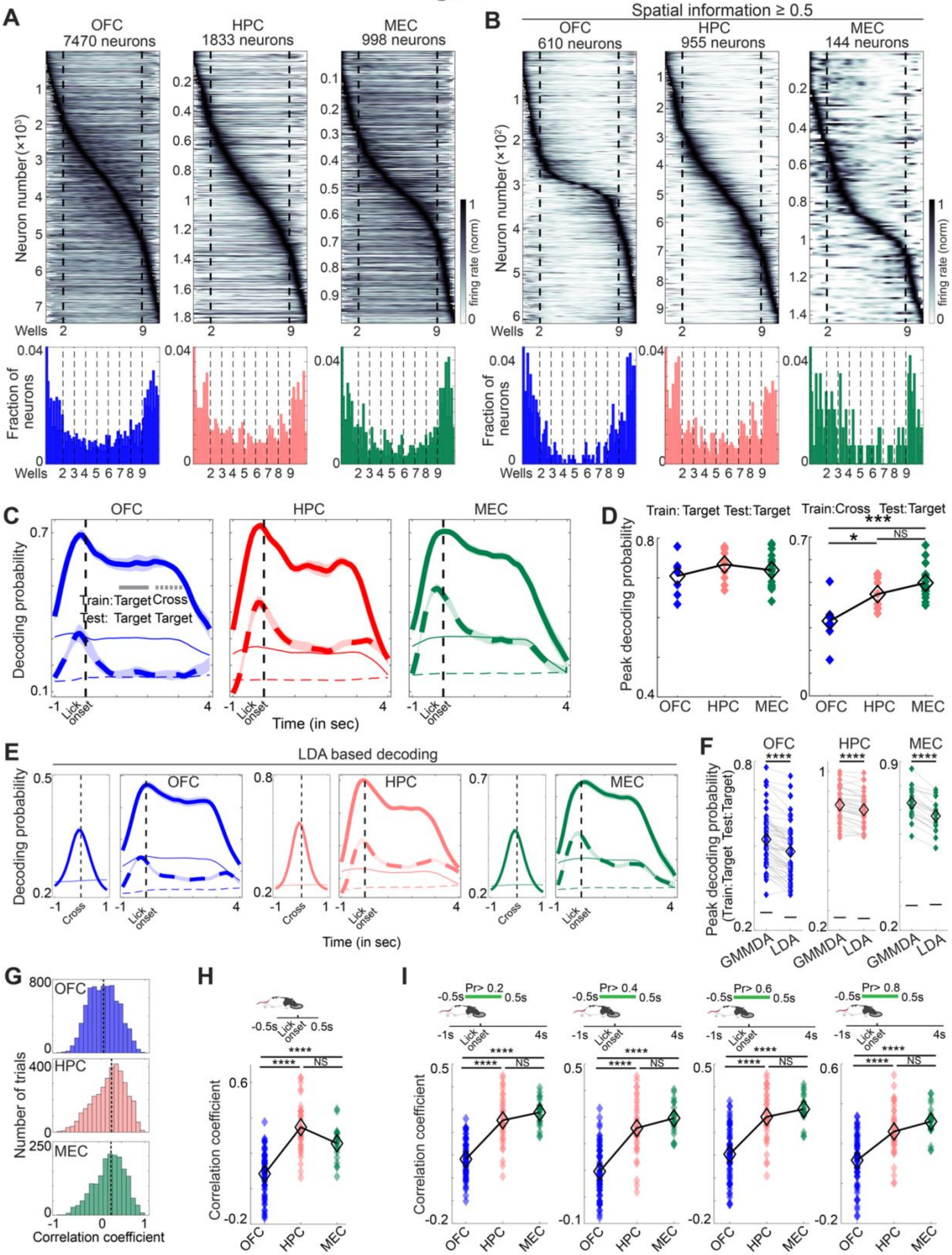
A
OFC 7470 neurons
HPC 1833 neurons
MEC 998 neurons
Neuron number (×10³)
firing rate (norm)
Wells
Fraction of neurons
B
Spatial information ≥ 0.5
OFC 610 neurons
HPC 955 neurons
MEC 144 neurons
Neuron number (×10²)
C
OFC
HPC
MEC
Decoding probability
Train: Target Cross
Test: Target Target
Lick onset
Time (in sec)
D
Train: Target Test: Target
Train: Cross Test: Target
Peak decoding probability
E
LDA based decoding
Cross
F
Peak decoding probability (Train: Target Test: Target)
GMMDA
LDA
G
Number of trials
Correlation coefficient
H
I
Pr> 0.2
Pr> 0.4
Pr> 0.6
Pr> 0.8
NS

24

**Figure S2: Spatial coding properties across brain regions (related to Figure 1)**: **(A)** Spatial firing rates of individual OFC, HPC, and MEC neurons, normalized to the peak firing rate and ordered based on the location of peak firing. The locations of wells 2 and 9 are marked by vertical dotted lines. For each neuron, the firing rate in the dominant running direction (defined as the direction with the higher peak firing rate) is plotted. Bottom row shows the distribution of the peak firing location of the neurons along the linear maze. Note that, corroborating previous observations, HPC neurons overrepresent the reward locations. Only neurons with peak firing rates above 1 Hz are shown. **(B)** Same as in A, except for neurons with spatial information (SI) exceeding 0.5. Note that only 8.17% of OFC neurons, in contrast to 52.1% of HPC neurons, exhibit an SI > 0.5. In addition, the peak firing locations of the spatially tuned OFC neurons are largely clustered around both ends of the linear maze, while those in the HPC neurons are more evenly distributed along the maze. These observations corroborate the differences in decoding the animal's instantaneous position across the brain regions (Figure 1D). Although only 14.43% of MEC neurons exceed an SI of 0.5, the high accuracy of the animal's current location coding in this region can be attributed to neurons with broad but consistent spatial tuning, such as putative grid cells with multiple firing fields or neurons whose firing rates ramp reliably along the maze. **(C)** Comparisons of target-well decoding probabilities using either the target-well decoder (solid lines) or the cross-well decoder (dashed lines) on a subset of sessions with overall similar decoding performance by the target-well decoder across the regions (n = 8 sessions in OFC, 10 sessions in HPC, and 17 sessions in MEC). Note the relatively weak target well decoding in the OFC using the cross decoder. **(D)** Quantification of the peak decoding probabilities in the plots shown in C. Left panel shows the similar decoding performance using the target-well decoder across the brain regions (means ± s.e.m.: 0.705 ± 0.016 in OFC; 0.734 ± 0.013 in HPC; 0.719 ± 0.01 in MEC; $p > 0.05$ for all pairwise comparison following Kruskal-Wallis test). Right panel indicates target-well decoding performance with the cross-well decoder was significantly lower in OFC than HPC or MEC (means ± s.e.m.: 0.33 ± 0.034 in OFC; 0.448 ± 0.022 in HPC; 0.498 ± 0.018 in MEC; $p=0.046$ for OFC versus HPC; $p = 0.00057$ for OFC versus MEC; $p = 0.465$ for HPC versus MEC by post-hoc pairwise comparison following Kruskal-Wallis test). **(E)** Decoding probabilities for non-target wells using the cross-well decoder (solid line on left, -1 to 1-second relative to crossing) and those for target wells using either the cross-well decoder (thick dotted line on right) or the target-well decoder (solid line on right, -1 to 4 seconds relative to lick onset). Decoding was performed on neural ensembles from OFC (left), HPC (middle), and MEC (right), using a Linear Discriminant Analysis (LDA)-based decoder. Analogous to Figure 1F. **(F)** Comparison of peak target-well decoding performance between GMMDA-based versus LDA-based target-well decoders. Horizontal lines denote the peak chance levels over the entire decoding window (-1s to 4s relative to lick onset). Note that the GMMDA-based decoder outperforms LDA-based decoders, although the latter still achieved performance significantly above chance levels. ****$p < 0.0001$ by Wilcoxon signed-rank test. **(G)** Histogram of the correlation coefficients between decoding probabilities obtained from the cross- and target-well decoders for all correct trials ($n = 7171$ trials for OFC; 3615 trials for HPC; 1730 trials for MEC). Dotted vertical lines refer to the medians. **(H)** Correlation coefficients between decoding probabilities obtained from the cross- and target-well decoders, computed over -0.5 s to 0.5 s relative to lick onset. **(I)** Correlation coefficients between decoding probabilities obtained from the cross- and target-well decoders, computed for trials with their mean target-well decoding probability during -0.5 s to 0.5 s relative to lick onset exceeding cutoff thresholds of 0.2, 0.3, 0.6, or 0.8 (panels from left to right). This analysis suggests that the difference in the OFC spatial coding between target- and cross-wells (Figure 1I) persists even when focusing on trials with high target-well decoding performance. Plotting scheme for panels S2H and S2I is the same as in Figure 1I. ****$p < 0.0001$ by post-hoc comparison following Kruskal-Wallis test.

#### Figure S3

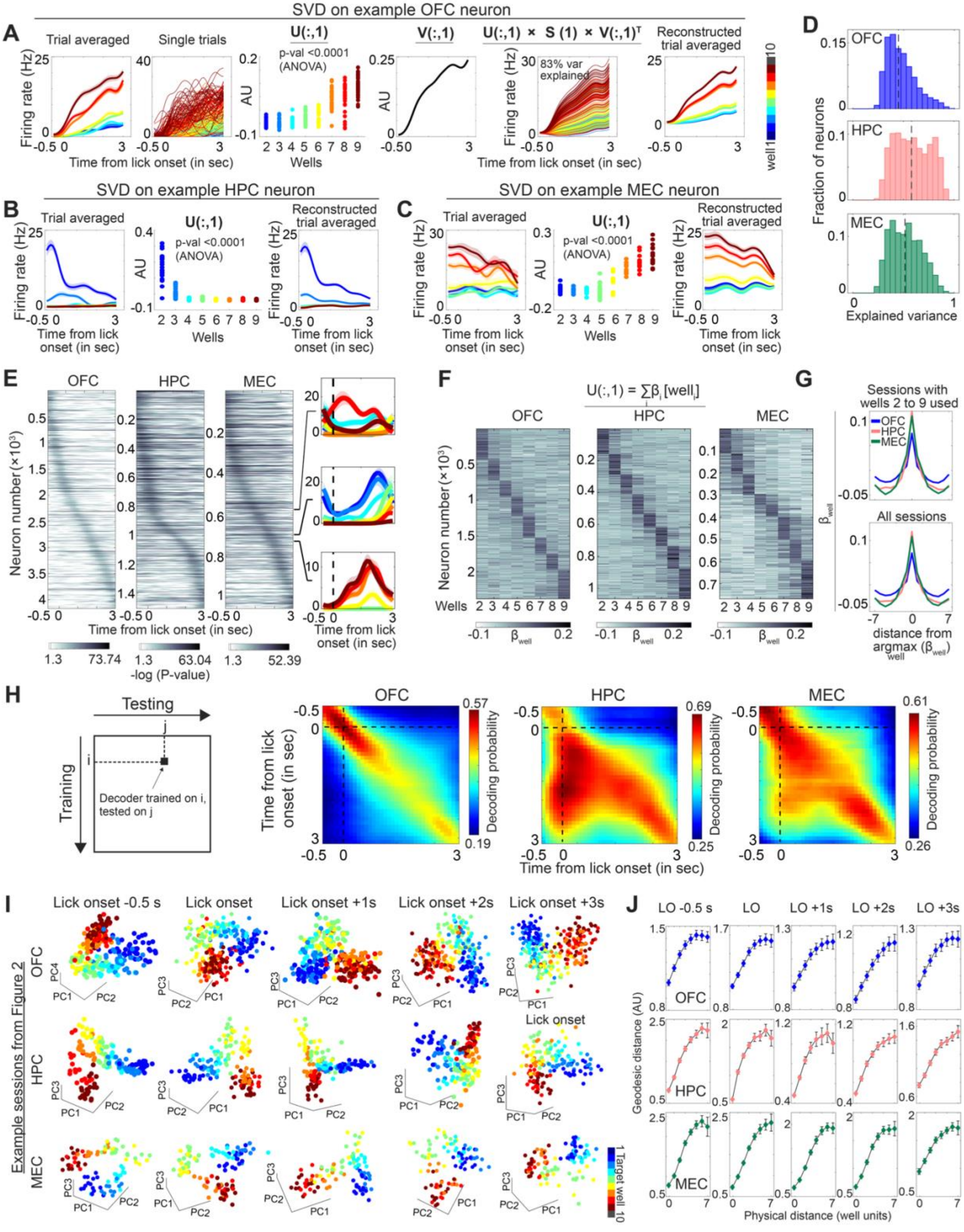
SVD on example OFC neuron
A
Trial averaged
Single trials
U(:,1)
p-val <0.0001 (ANOVA)
V(:,1)
U(:,1) × S (1) × V(:,1)T
83% var explained
Reconstructed trial averaged
Firing rate (Hz)
Time from lick onset (in sec)
Wells
AU
well 1
10
D
OFC
HPC
MEC
Fraction of neurons
Explained variance
SVD on example HPC neuron
B
SVD on example MEC neuron
C
Time from lick onset (in sec)
E
OFC
HPC
MEC
Neuron number (×10³)
-log (P-value)
F
U(:,1) = ∑βi [welli]
Wells
βwell
G
Sessions with wells 2 to 9 used
All sessions
distance from argmax (βwell)
H
Testing
Training
Decoder trained on i, tested on j
Time from lick onset (in sec)
Decoding probability
I
Lick onset -0.5 s
Lick onset
Lick onset +1s
Lick onset +2s
Lick onset +3s
Example sessions from Figure 2
Target well
J
LO -0.5 s
LO
LO +1s
LO +2s
LO +3s
Geodesic distance (AU)
Physical distance (well units)

76

**Figure S3: Dynamic and well-specific coding properties of OFC, HPC, and MEC neurons (related to Figure 2)**: **(A)** Example of singular value decomposition (SVD) based extraction of single-trial contributions for an example OFC neuron. First two panels from the left show the trial-averaged and single-trial firing patterns aligned to lick onset. SVD-based factorization of single trial activity results in an orthonormal matrix U, whose columns represent the unique contributions of individual trials (third panel), an orthonormal matrix V, whose columns abstract the temporal dynamics within the trial (fourth panel), and a diagonal matrix S whose elements represent the amount by which the outer products of U and V should be scaled. The fifth and sixth panels depict the reconstructed single trial and trial-averaged firing rates from just the first columns of U and V, which explains 83% variance of the original single trial firing rate matrix. Note the similarity in the original and reconstructed firing rates. In this paper, the first column of U is referred to as the single-trial contribution of a neuron. **(B-C)** SVD-based single-trial contributions extracted for representative HPC (B) and MEC (C) neurons. **(D)** Distribution of the fractions of firing-rate variances explained by the single-trial contributions (i.e., the first column of U) for individual well-specific neurons from each brain area. Dashed vertical lines represent the median (0.449 for OFC, 0.579 for HPC, and 0.516 for MEC). **(E)** Statistical significance of well-position coding of individual neurons during approach and licking. Neurons with significant well-specific activity from each brain region are arranged according to their peak well discriminability relative to lick onset. For each neuron, the firing rate time series was segmented into 100-ms bins, followed by an ANOVA performed at each time bin. The ANOVA-based p-values are corrected for multiple comparisons using Bonferroni corrections. Note that OFC neurons, in general, discriminate well positions (with significant p-values) at smaller time windows compared to HPC and MEC neurons, suggesting that, at an ensemble level, the OFC spatial representation is highly dynamic. Insets on the right show the trial-averaged firing rates of three MEC neurons that exhibit well-specific activity largely during licking compared to the target well approach phase. **(F)** Between-time-point decoding of target-well identities. The schematic on the left outlines the analysis procedure where a decoder is trained on the neural activity at a given time point and tested at another time point. Panels 2 to 4 from left show the between-time-point decoding from each brain region, averaged across sessions. Consistent with panel E, target-well decoding in OFC neural ensembles is highly dynamic, such that a decoder trained at a given time point does not generalize to other time points, resulting in high decoding mainly along the diagonal. In contrast, HPC and MEC neural ensembles exhibit a lower degree of time dependency, resulting in a more generalizable spatial code during the lick duration. **(G)** Coding strength $\beta_{well}$ of individual neurons, defined as the well-specific regression coefficients explaining single-trial contributions U(:,1). Neurons are arranged based on their peak well coding strengths. Shown are well-specific neurons from sessions where all eight wells from well 2 to well 9 were used. **(H)** Individual $\beta_{well}$ values, plotted against their distances from the well with the maximum $\beta_{well}$, averaged over all neurons shown in panel G (top) and well-specific neurons from all sessions (bottom). Note that the $\beta_{well}$ values in OFC neurons reduce more gradually with distance from their peaks, compared to HPC and MEC neurons, suggesting that individual OFC neurons encode nearby wells with similar firing rates. **(I)** Neural ensemble activity at five different time points relative to lick onset projected onto three PC axes. Same example sessions as those shown in Figures 2D, 2H, and 2I. Note that the ensemble neural activity at individual time points reveals the topological arrangement according to well positions (similar to those in Figure 2). **(J)** Geodesic distances between individual target-well representations, computed over neural ensemble activity at different time points relative to lick onset (LO), plotted against the physical distances between the corresponding wells (as in Figure 2J). To match the number of TC dimensions used in Figure 2, the geodesic distances were computed over $n = 8$, 6, and 6 PC dimensions in OFC, HPC, and MEC sessions, respectively. Diamonds and error bars represent means and s.e.m. across $n = 62$ sessions in OFC, 37 sessions in HPC, and 21 sessions in MEC. Note the near-linear relationships between neural distances and physical distances in all three brain areas at all the time points. Overall, panels I and J reveal that the topology-preserved spatial coding revealed by TCA (Figure 2) is also evident in neural ensemble activity at different lick-duration time points. However, due to dynamic coding, especially in OFC, abstracting temporal components using TCA (or SVD, as shown in panel A) circumvents the need to choose a specific time point.

#### Figure S4

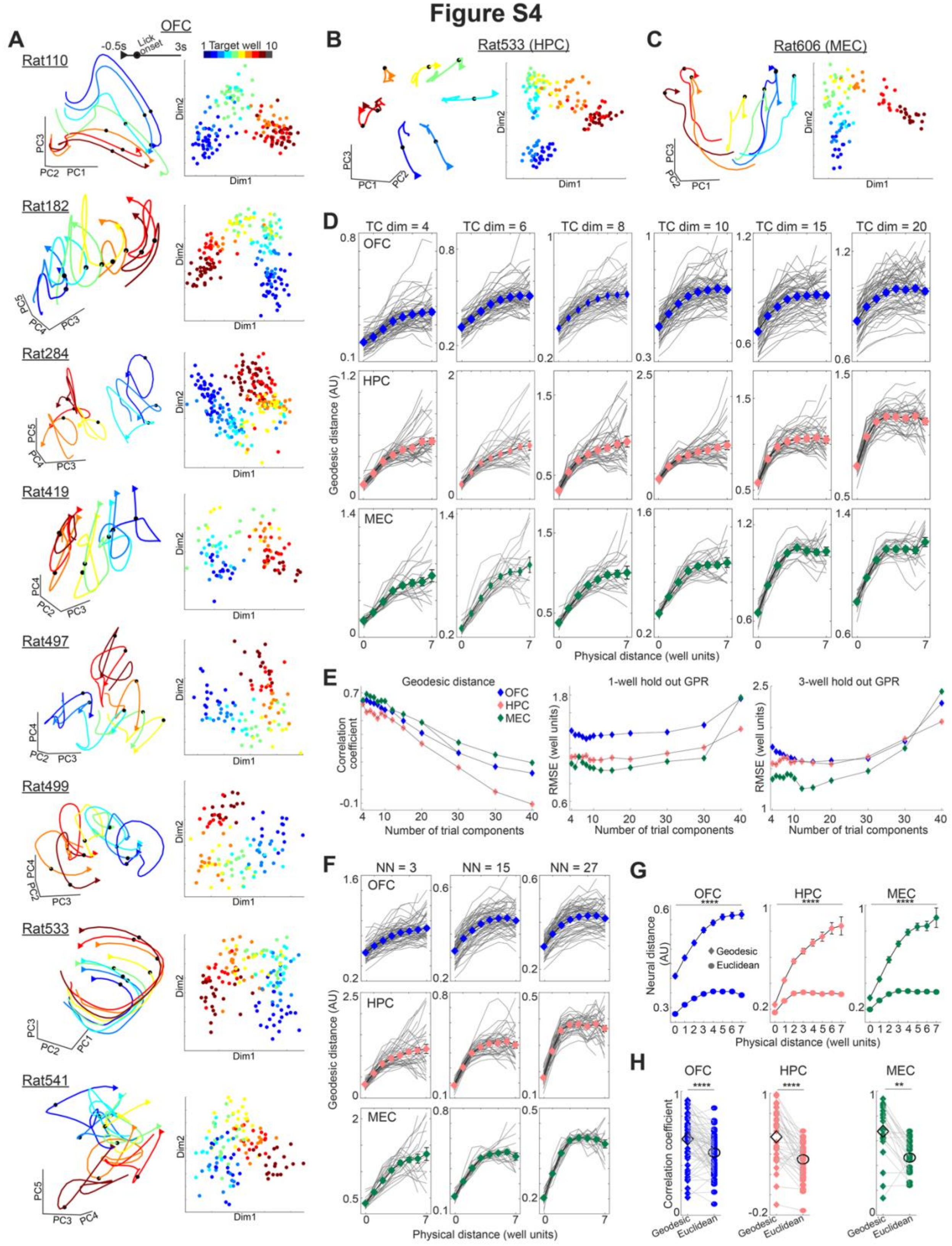


133

**Figure S4: Topology-preserved spatial mapping across animals and analysis conditions (related to Figure 2): (A)** (left) Trial-averaged OFC neural ensemble activity from representative sessions from eight rats projected on three PC axes. Neural activity from 0.5 sec before to 3 sec after the lick onset is shown. The right panels depict the corresponding first eight trial components (from TCA) projected on two LDA dimensions (orthogonalized) that maximize the separation between target-well identities. Note that the topological mapping of well location is visible in the OFC of all recorded animals. **(B-C)** Same as A but for neural ensembles recorded from HPC (B) and MEC (C). Panels A-C present sessions distinct from the ones plotted in Figures 2B, 2D, 2H, and 2I. **(D)** Neural geodesic distances between target-well representations as a function of the physical distances between the corresponding wells, computed for varying numbers of trial components (TC dimensions). The near-linear relationships between geodesic and physical distances persist at low numbers of trial components, although increasing the dimensionality (for example, the last two columns from right) leads to more uniform distance distributions, resulting in a plateau of geodesic distances over larger physical distances. In Figure 2, eight dimensions for OFC and six dimensions for HPC and MEC were used. **(E)** (left) Correlation coefficients between geodesic neural representational distances and physical well distances, averaged across sessions and plotted as a function of the number of trial components used. Note the relatively high correlation coefficients at low dimensions. Middle and right panels show session-averaged root-mean-squared errors (RMSEs) in location predictions obtained using either 1- or 3-well held-out GPR analyses performed on increasing numbers of trial components. GPR-based prediction is less sensitive to dimensionality and only fails at very high dimensions. **(F)** Neural geodesic distances between target-well representations obtained by using different nearest neighbor (NN) values to compute the adjacency graph for the modified Isomap algorithm. NNs reflect the scale at which a manifold is analyzed. A low number of NN results in a very local view of the manifold, and even nearby points may be excluded from being the nearest neighbor, while high NN values can overlook the curvature of the manifold by grouping distal points as nearest neighbors (note the plateauing of geodesic distances in the middle and right panels). Hence, we chose an intermediate value of NN = 6. **(G-H)** Geodesic (diamond) versus Euclidean (circle) distances between target-well representations (G) and their correlations with the physical distances between the corresponding wells (H), from all three brain regions. Corroborating the curved nature of the manifolds, Euclidean measures of neural distance are significantly smaller than their geodesic counterparts at all physical distances (by Wilcoxon signed-rank test). In addition, the reduced correlation coefficients (panel H) suggest that geodesic distances better reflect the spatial arrangement of physical well positions. ****$p < 0.0001$, **$p < 0.01$ by Wilcoxon sign-rank test. In panel S4G, diamonds and circles represent the means, while error bars represent the s.e.m. over sessions. In panel S4H, solid and open symbols depict individual sessions and their means, respectively. Plotting scheme in panels S4D and S4F is identical to that in Figure 2J.

### Figure S5

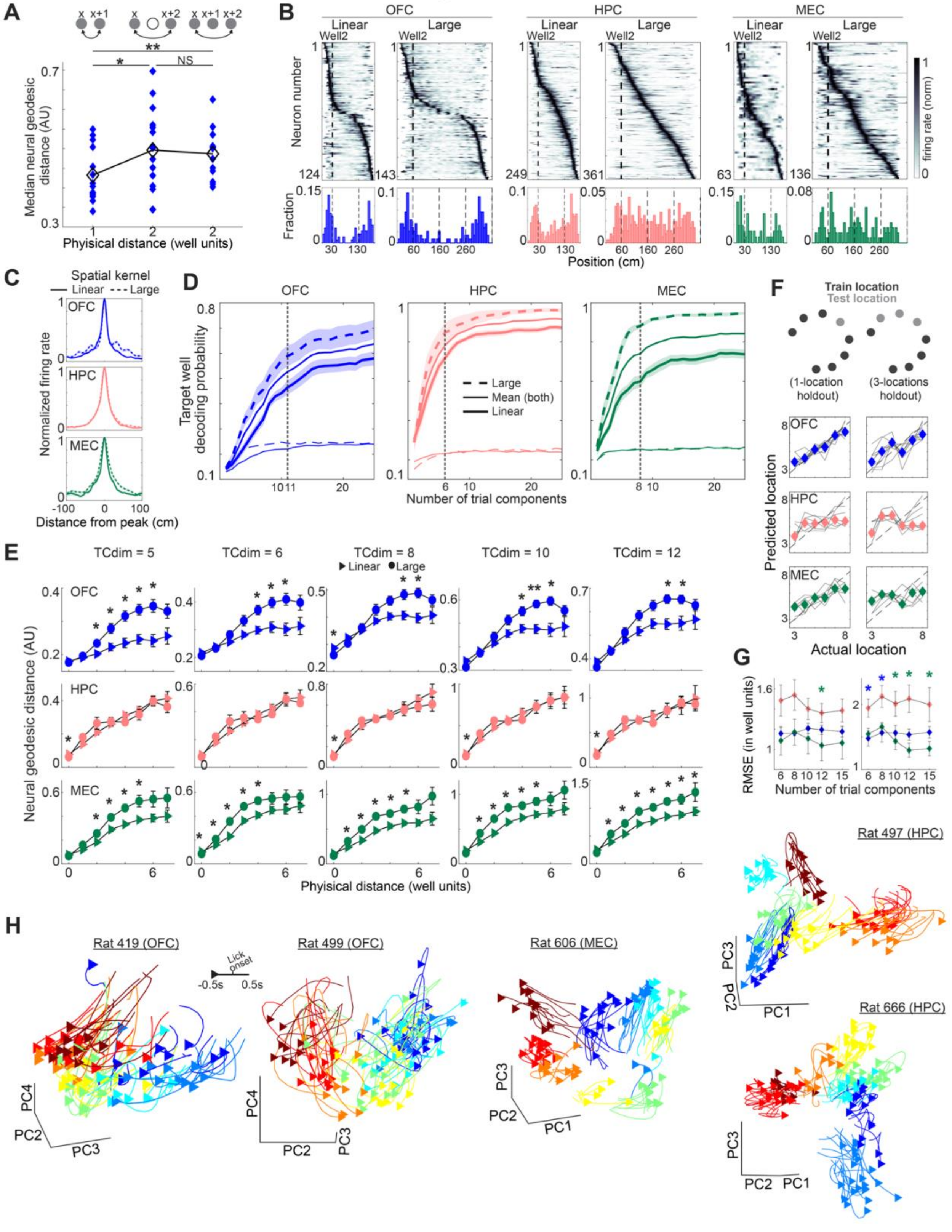

A
x x+1
x x+2
x x+1 x+2
**
*
NS
Median neural geodesic distance (AU)
Phyisical distance (well units)
B
OFC
HPC
MEC
Linear
Large
Well2
Neuron number
firing rate (norm)
Fraction
Position (cm)
C
Spatial kernel
Linear
Large
OFC
HPC
MEC
Normalized firing rate
Distance from peak (cm)
D
OFC
HPC
MEC
Target well decoding probability
Large
Mean (both)
Linear
Number of trial components
F
Train location
Test location
(1-location holdout)
(3-locations holdout)
Predicted location
Actual location
E
TCdim = 5
TCdim = 6
TCdim = 8
TCdim = 10
TCdim = 12
Linear
Large
Neural geodesic distance (AU)
Phyisical distance (well units)
G
RMSE (in well units)
Number of trial components
H
Rat 419 (OFC)
Rat 499 (OFC)
Rat 606 (MEC)
Rat 497 (HPC)
Rat 666 (HPC)
Lick onset
-0.5s
0.5s
PC1
PC2
PC3
PC4

**Figure S5: OFC spatial map reflects not only topology but also distance of physical well locations (related to Figure 3). (A)** OFC neural geodesic distances computed between representations of either adjacent wells, wells separated by an unused intermediate well, or wells separated by a used intermediate well. OFC neurons preserve the distance between wells irrespective of whether the well in between was rewarded or not during the session, implying that OFC does not just preserve the order of target well locations, but also their distances. Colored diamonds and black open diamonds represent individual sessions (n = 14 sessions) and their means, respectively. **(B)** Spatial firing rates of individual OFC, HPC, and MEC neurons in both linear and large mazes, normalized to the peak firing rates and ordered based on the locations of peak firing. Only neurons with SI > 0.5 and peak firing > 1 Hz are plotted. Firing rates are aligned to the coordinate of well 2. Bottom row shows the distribution of the peak firing location of the neurons along the tracks. The firing fields of spatially-tuned OFC neurons are distributed near the ends of both mazes, while HPC and MEC neurons tile across the mazes. **(C)** Spatial kernels of the neurons shown in panel B above, computed as the firing rates over a 201 cm stretch centered at the peak firing location. Note the similarity in the kernels between the mazes, especially for HPC neurons, implying largely similar place cell properties in the large maze compared to the linear maze. **(D)** Target-well decoding probabilities in the linear and the large mazes for each brain region. Solid lines show the decoding probabilities averaged across the two sessions. Dotted vertical lines represent the number of TCA trial components used for computing the neural geodesic distance in Figure 4F. n = 11, 6, and 8 components explained 82.77%, 85.43%, and 87.88 % of the average decoding levels observed when using 25 trial components. **(E)** Geodesic distances computed using varying numbers of TCA trial components from OFC (top), HPC (middle), and MEC (bottom) ensembles plotted against the physical distances (in well units) for both linear and large mazes. Note that the well units in the large maze are twice as large as those in the linear maze. Shown are the means (filled triangle and circle) ± s.e.m. (errorbars). *$p < 0.05$, **$p < 0.01$ by Wilcoxon sign-rank test. The increased scale of the spatial mapping in the large maze can be observed along a wide range of TCA trial-component numbers obtained from OFC and MEC neural ensembles. Similarly, the lack of scaling in HPC neural ensembles is also independent of the number of TCA trial components used. **(F)** GPR-based prediction of target-well identity in the large maze from OFC, HPC, and MEC trial components. Left panel shows predictions when a single well was held out. Right panel shows a prediction strategy where wells 3, 4, and 5, or wells

6, 7, and 8, were held out. Each gray curve represents the median predictions from a single session, while the colored diamonds depict the medians across sessions. N = 11, 6, and 8 trial components were used for this analysis. **(G)** Root mean squared errors (RMSEs) of well predictions with means ± s.e.m. for the single- (left) and three (right)-well holdout strategies computed for varying numbers of TCA trial components. Overall, the GPR-based well predictions are worse in HPC. Blue and green stars indicate $p < 0.05$ for post hoc OFC versus HPC and MEC versus HPC, respectively, following Kruskal-Wallis test. **(H)** Plots showing ensemble neural activity in OFC, MEC, and HPC from example sessions in the large maze. Trial-wise neural activity, color-coded by target well identity, is projected on three principal components. For each trial, one second of activity centered at lick onset is shown. Note the topology-preserved mapping of well locations in the OFC and MEC neural ensembles, but less so in the HPC neural ensembles.

### Figure S6

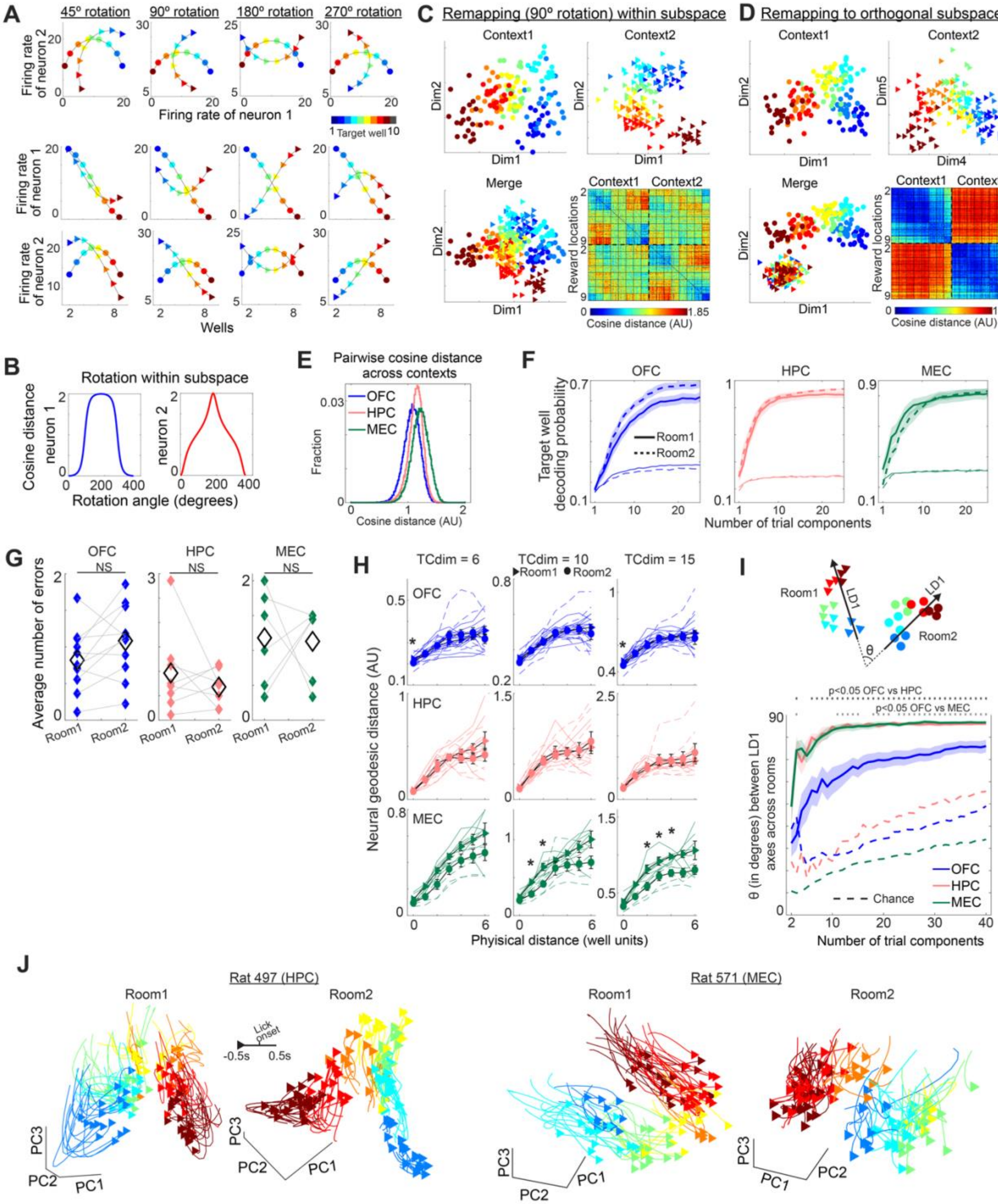
A
45º rotation
90º rotation
180º rotation
270º rotation
Firing rate of neuron 2
Firing rate of neuron 1
Target well
Firing rate of neuron 1
Firing rate of neuron 2
Wells
B
Rotation within subspace
Cosine distance neuron 1
neuron 2
Rotation angle (degrees)
C
Remapping (90º rotation) within subspace
Context1
Context2
Merge
Dim1
Dim2
Reward locations
Cosine distance (AU)
D
Remapping to orthogonal subspace
Context1
Context2
Merge
Dim1
Dim2
Dim4
Dim5
Reward locations
Cosine distance (AU)
E
Pairwise cosine distance across contexts
OFC
HPC
MEC
Fraction
Cosine distance (AU)
F
OFC
HPC
MEC
Target well decoding probability
Room1
Room2
Number of trial components
G
OFC
HPC
MEC
NS
Average number of errors
Room1
Room2
H
TCdim = 6
TCdim = 10
TCdim = 15
Room1
Room2
OFC
HPC
MEC
Neural geodesic distance (AU)
Physical distance (well units)
I
Room1
Room2
LD1
θ
p<0.05 OFC vs HPC
p<0.05 OFC vs MEC
θ (in degrees) between LD1 axes across rooms
OFC
HPC
MEC
Chance
Number of trial components
J
Rat 497 (HPC)
Rat 571 (MEC)
Room1
Room2
Lick onset
-0.5s
0.5s
PC1
PC2
PC3

**Figure S6: Spatial mapping across different environments (related to Figure 4):** (A-D) Effect of remapping within position-coding subspaces (A-C) and to their corresponding orthogonal subspaces (D) using simulated activity. **(A)** The activity of two simulated neurons at eight different reward wells (colored) across two contexts (circle and triangle). The map in context 2 (triangle) is rotated at various angles (columns) relative to that in context 1 (circle) to resemble remapping within subspace (while maintaining the topological nature of spatial coding). Top row shows the maps in the neural activity space, where the horizontal and vertical axes depict the firing rates of neurons 1 and 2, respectively. Middle and bottom rows plot the location-specific firing rates of neuron 1 and neuron 2. Note the varying degree of similarity in well-specific firing across the two contexts as the maps rotate relative to each other. **(B)** Cosine distances between vectors representing the well-specific firing rates in each context plotted for neuron 1 (left) and neuron 2 (right) as the map in context 2 is rotated from 0 degrees to 359 degrees relative to the map in context 1. The highest cosine distance is achieved when one map is completely inverted relative to the other. For other rotation degrees, neurons exhibit varying degrees of cosine distances. **(C)** Simulated trial-wise spatial representations in the position-coding subspaces. To resemble high-dimensional noisy neural data, we implemented the following steps. The center of each reward well (8 wells used) representation was positioned along a semicircle of radius r with equal angular spacing. The 2-dimensional centers were then individually jittered along a third dimension by a random distance sampled from a standard normal distribution scaled by 0.2*r. This resulted in 3-dimensional coordinates for the well centers. Next, to simulate high dimensional single trial data, we converted the 3-dimensional coordinates into 20-dimensional by padding the remaining dimensions with zero and subsequently sampled 20 'trials', each 20-dimensional, for every well using a multivariate Gaussian distribution (using Matlab's mvnrnd function). For each well, the means of the Gaussian distributions comprised the 20-dimensional well centers. The covariance matrix was a 20-dimensional identity matrix multiplied by 0.5*r. Remapping within the subspace was performed by a rotation by 90 degrees on the plane of 2-dimensional well-centers. Subsequent steps were carried out independently for the two maps, meaning that the remapped well centers are not a replica of the original map (due to the independent jitter in the third dimension). The top two plots show the original map in context 1 (left) and the rotated map in context 2 (right). Bottom row plots the overlap of the two maps (left) and the trial-wise cosine distance matrices, as shown in the main Figures 4C and 6C. **(D)** Same as panel C, but the map in context 2 is projected onto an orthogonal set of dimensions. As the well centers in context 1 are 3-dimensional, the well centers in context 2 occupied dimensions 4 to 6 (with zeros padded in dimensions 1-3 and 7-20). Note the uniformly high cosine distances between trials in context 1 versus those in context 2, in contrast to the broadly distributed moderate distances when remapping occurs within the subspace (panel C). **(E)** Histogram of the cosine distances between trials across the two rooms, as shown in Figures 4C and 4D. Note that the cosine distances were more narrowly distributed in HPC compared to OFC and MEC (standard deviation: 0.143 for OFC; 0.126 for HPC; and 0.145 for MEC), while the overall distances were higher in both HPC and MEC compared to OFC (median: 1.077 for OFC; 1.146 for HPC; 1.217 for MEC). **(F)** Decoding probabilities of target well in each of the two rooms plotted against increasing number of trial components. Note that the decoding performance is largely similar between the rooms in all three brain regions. Solid line with shading indicates means ± s.e.m. across sessions. **(G)** Average number of errors (licks at incorrect wells) made by the animals per well-combination in individual rooms for each paired session (colored diamonds). Task performance was similar, in spite of a change in spatial context. Open diamonds represent the means. **(H)** Geodesic distances computed using varying number of TCA trial components from OFC (top), HPC (middle), and MEC (bottom), plotted against the physical distances (in well units) in each of the two rooms. Shown are the means (filled triangle and circle) ± s.e.m. (errorbars). *$p < 0.05$, by Wilcoxon signed-rank test. OFC and HPC spatial maps exhibit similar scaling in both rooms, while the MEC map in room 2 shows slight shrinkage. **(I)** The angles between the spatial maps from the two rooms computed in the TC space over an increasing number of trial components. For each map, LDA was performed in the common TC space to identify the LD1 dimension that explains the largest separation between the wells. For a given number of trial components, pairwise comparison for significance testing was performed following Kruskal-Wallis test. The angles between the maps are significantly smaller in OFC compared to those in HPC and MEC neural ensembles. As angles between vectors tend to increase with dimensionality, we computed the chance levels by randomly partitioning the trials from both sessions into two halves. Partitioning was performed 100 times, and the 95$^{th}$ percentile of the angles between the LD1 axes computed for each partitioned pair was considered as the chance level. Note that chance levels increase with the number of dimensions, but the angle between LD1 axes is above the chance levels in all three brain areas (except dimensions 2-4 for OFC), implying that the observed angles between the two context-dependent maps are not merely a result of two random high-dimensional vectors. **(J)** Plots of neural ensemble activity in HPC and MEC from a representative paired session each, projected onto the first three PCs. The PCs are computed separately for each room to highlight that, in spite of global remapping, topology-preserved maps are formed independently in each room.

### Figure S7

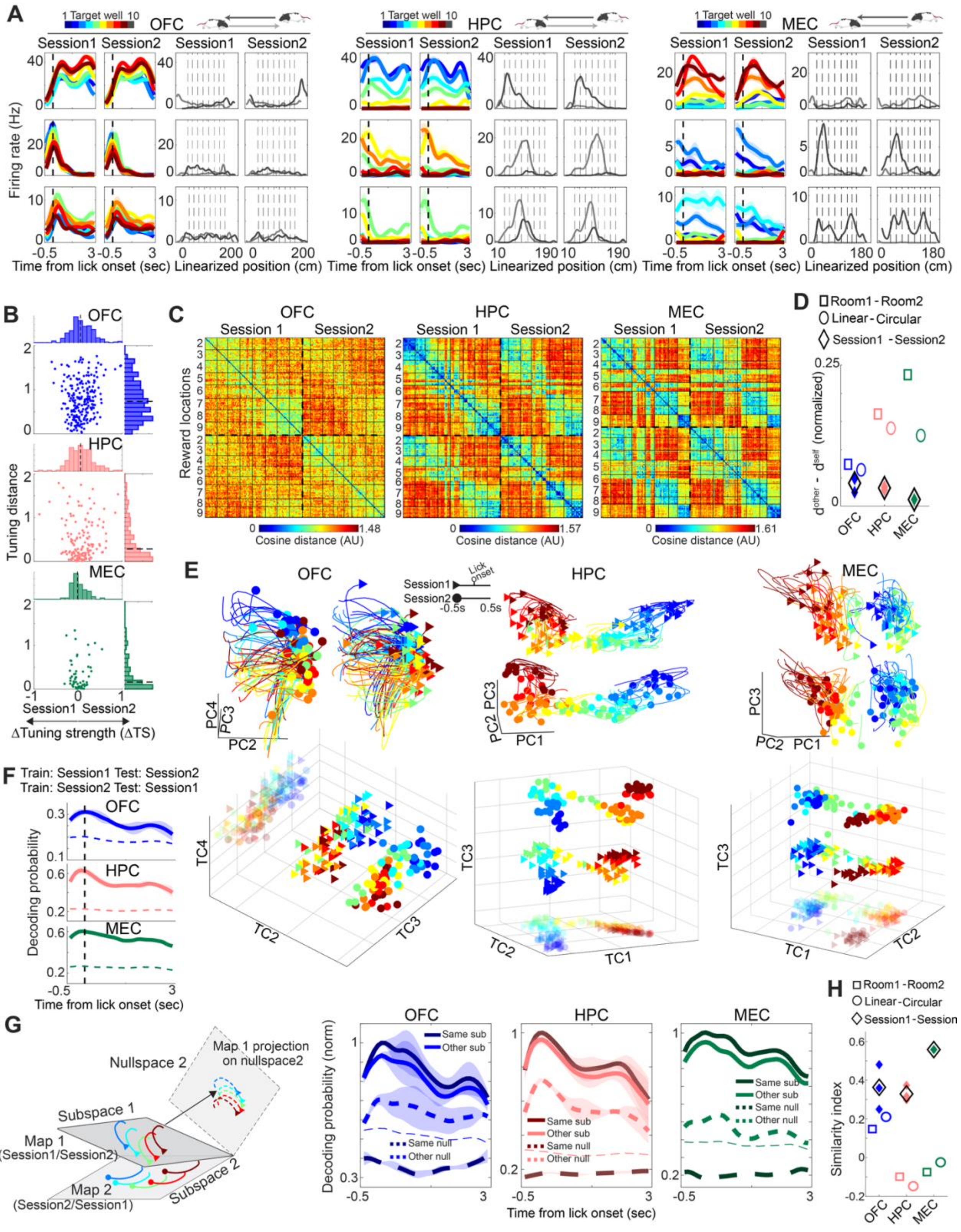


284

**Figure S7: OFC, HPC, MEC maintain stable spatial maps within the same context (related to Figures 4 and 6): (A)** Firing rates of three representative neurons from OFC (left), HPC (middle), and MEC (right) in the linear maze between two consecutive sessions recorded in the same room (n = 3 sessions from 3 rats for OFC; 3 sessions from 3 rats for HPC; 1 session from 1 rat for MEC). For each neuron, the two left panels show well-specific firing rates aligned to lick onset, whereas the right panels display firing rates across maze positions. Activity for the two running directions is plotted in separate shades (light and dark grey). Dotted vertical lines represent well locations (wells 2-9 were used). **(B)** Tuning distances versus ΔTS, plotted for OFC (top), HPC (middle), and MEC (bottom) neurons. Each dot corresponds to a single neuron. Marginal distributions are plotted on the top and right of each panel. Dotted lines represent the medians. Note that, unlike Figures 4B and 6B (which show data across different rooms and different maze geometries, respectively), the ΔTS and tuning distance of HPC and MEC neurons are distributed close to zero. **(C)** Distance matrices of trial-by-trial ensemble neural representations for a representative paired session from OFC (left), HPC (middle), and MEC (right). Each pixel depicts the cosine distance between the population vectors representing two individual trials. Trials are grouped according to the target-well identity, and trials from the two sessions are separated by thick dotted lines. In all three brain regions, the top left and top right quadrants share a similar cosine distance pattern, signifying the similarity in neural population coding between the two sessions. **(D)** The differences between $d^{other}$ and $d^{self}$, normalized to their sums and averaged across all well pairs in individual sessions. The values of $d^{other}$ and $d^{self}$ are computed as described in the schematics of Figures 4D and 6D. Filled colored and open black diamonds represent individual sessions and their means, respectively. For comparison, the average values from sessions recorded in different rooms (open rectangles, values same as those in Figure 4D) and across different maze geometries (open oval, values same as those in Figure 6D) are plotted. Unlike OFC, HPC and MEC neurons exhibit higher $d^{other}$ - $d^{self}$ values when either the room or the maze geometry is changed. **(E)** Plots of ensemble neural activity in OFC (left), HPC (middle), and MEC (right) in the two sessions from representative paired sessions. Top panel plots trial-wise neural activity, color-coded by the target-well identity and projected on three principal components. Bottom panels show three TCA trial components of the corresponding sessions. Translucent triangles and circles are projections of the trial components on a single plane. To facilitate comparisons of neural representations between the two sessions, the representations from one session are shifted along a single dimension. Spatial mapping of the well locations is almost identical between consecutive sessions in all three brain regions. **(F)** Decoding probabilities of target well in one of the two sessions using a decoder trained on the other. Thin dotted lines show the chance levels. Shown are means (solid) ± s.e.m. (shading). Note the higher decoding probability in HPC and MEC ensembles compared to that observed across different spatial contexts (Figures 4F and 6F) **(G)** (Left) Schematic of the subspace-based decoding analysis. (Right) Decoding probabilities of target well from neural activity projected on the distinct subspaces occupied by the spatial maps from the two sessions, as well as their corresponding null spaces. Shown are the means (solid and thick dotted lines) ± s.e.m. (shading). Thin lines show the chance levels for decoding on the same subspaces. For each paired session, the decoding probabilities are normalized to the decoding probability on the same subspace at lick onset. **(H)** Subspace similarity index (see main text), plotted for each paired session (individual colored diamonds) from all three brain areas. Open black diamonds depict the means. For comparison, the average values from sessions recorded in different rooms (open squares, values same as those in Figure 4G) and in mazes with different geometries (open circles, values same as those in Figure 6G) are plotted. Note that the maps across two consecutive sessions reside in similar subspaces in all three brain regions, resulting in high subspace similarity indices.

### Figure S8

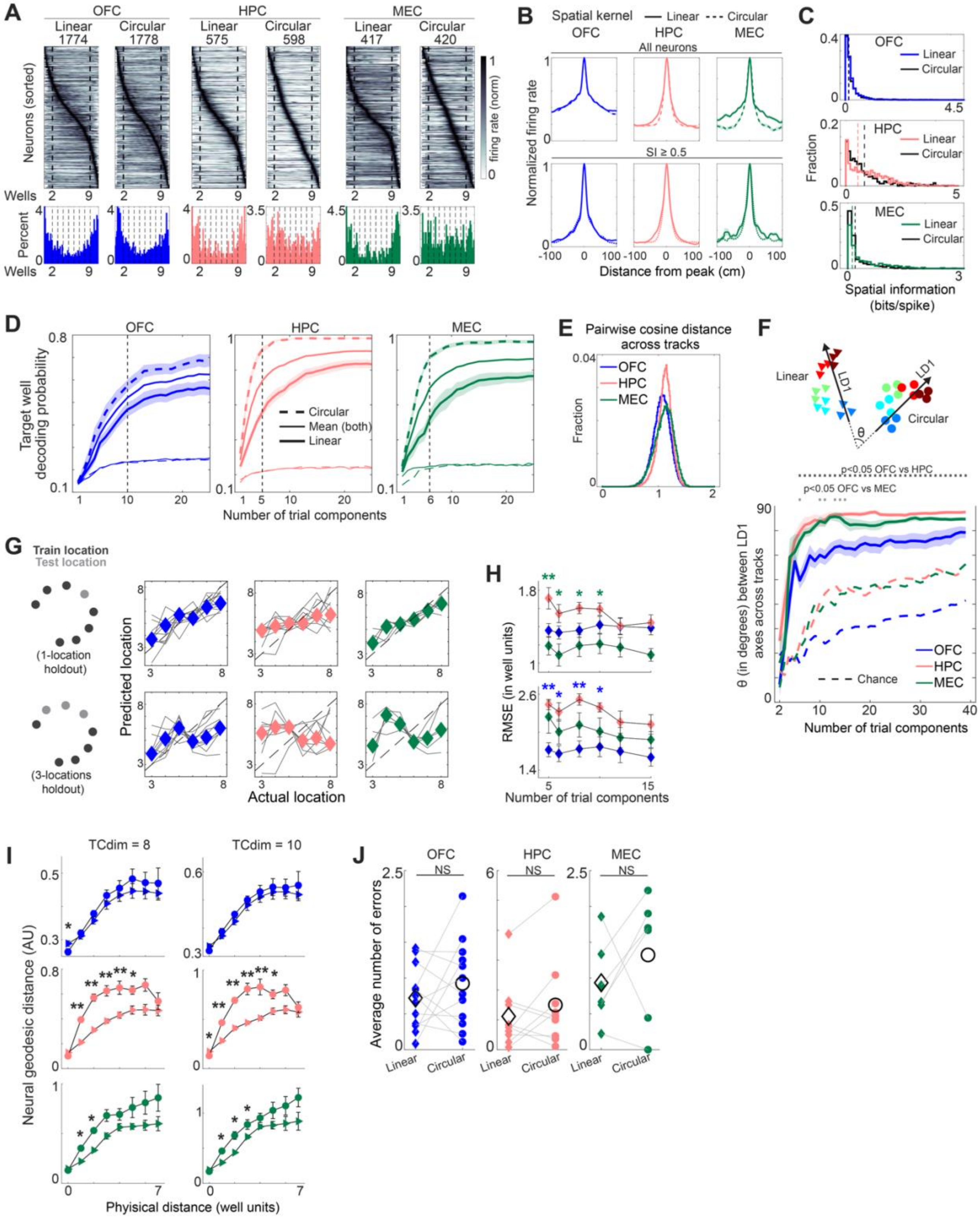


333

334

**Figure S8: Spatial mapping across different maze geometries (related to Figures 5 and 6): (A)** Spatial firing rates of individual OFC, HPC, and MEC neurons in both linear and circular mazes, normalized to the peak firing rates and ordered based on the peak-firing locations. All neurons with peak firing > 1 Hz are plotted. Locations of wells 2 and 9 are marked with vertical lines. Bottom row shows the distributions of the peak firing location of the neurons along the mazes. The firing field distributions across mazes are almost identical for OFC neurons. However, HPC and MEC neurons exhibit a slight accumulation of firing fields near the ends in the linear maze, while they uniformly tile the circular maze. Numbers on top show the number of neurons. The y-axis of the histograms for the linear maze has been truncated due to a high number of neurons occupying the leftmost bin. **(B)** Spatial kernels, computed as the firing rates over a 201 cm stretch centered at the peak firing locations. Top row includes all neurons with peak firing rates above 1 Hz, while the bottom row depicts the results from neurons with spatial information greater than 0.5. Note that the spatial kernels in linear and circular mazes are identical for OFC neurons, while those for HPC and MEC neurons are narrower in the circular maze, implying the differences in spatial tuning widths. **(C)** Distributions of spatial information for the neurons shown in panel A. Corroborating the spatial kernel results, spatial information distribution is identical between the two mazes for OFC neurons, but significantly right shifted for HPC and MEC neurons in the circular maze (median SI linear and circular: 0.128 and 0.135 for OFC; 0.538 and 0.829 for HPC; 0.117 and 0.191 for MEC; $p = 0.419$, $1.163 \times 10^{-8}$, and $1.181 \times 10^{-6}$ for linear versus circular by Kolmogorov-Smirnoff test for OFC, HPC, and MEC, respectively). **(D)** Decoding probabilities of target well in the linear and the circular mazes for each brain region. Solid lines show the decoding probabilities averaged across the two sessions. Dotted vertical line represents the number of TCA trial components used for computing the neural geodesic distances in Figure 5D. n = 10, 5, and 6 dimensions explain 83.51%, 81.28%, and 80.62 % of the average decoding levels obtained when using 25 trial components. The significantly higher decoding performance observed in the circular maze with HPC and MEC trial components can in part be attributed to the high spatial information and sharper spatial kernels of neurons in these brain regions (panels B and C). **(E)** Histogram of the cosine distances between trials across the two mazes, as shown in Figures 6C and 6D. Note that the cosine distances are more narrowly distributed in HPC compared to OFC and MEC (standard deviation: 0.147 for OFC; 0.128 for HPC; and 0.168 for MEC), while the overall distance was higher in both HPC and MEC compared to OFC (median: 1.066 for OFC; 1.12 for HPC; 1.115

for MEC). **(F)** Angles between the spatial maps for the two mazes computed in the TC space over an increasing number of trial components. Plotting and analysis are similar to Figure S6-I. *$p < 0.05$, **$p < 0.01$, pairwise comparison for significance testing was performed following Kruskal-Wallis test. The angles between the maps were significantly smaller in OFC compared to those in HPC and MEC, especially at lower dimensions. **(G)** GPR-based predictions of target-well identity in the circular maze from OFC, HPC, and MEC trial factors. Top row shows predictions when a single well was held out. Bottom row shows a prediction strategy where either wells 3, 4, and 5, or wells 6, 7, and 8, were held out. Each gray curve represents the median predictions from a single session, while the colored diamonds depict the medians across sessions. When three wells are held out, location prediction is largely impaired in HPC. Trial components of $n = 10$, 5, and 6 were used for this analysis. Note that, based on the results of decoding performance in panel D, these numbers of components are sufficient for decoding well identities in all three brain regions. **(H)** Root mean square errors (RMSEs) of well prediction, with means ± s.e.m. (colored diamonds and error bars), for either the single- (top) or three- (bottom) well holdout strategies computed for varying numbers of TCA trial components. Overall, the GPR-based well prediction is worse in HPC. Blue and green stars indicate significance (*$p < 0.05$, **$p < 0.01$) for post hoc OFC versus HPC and MEC versus HPC, respectively, following Kruskal-Wallis test. **(I)** Geodesic distances of neural representations computed using varying number of TCA trial components from OFC (top), HPC (middle), and MEC (bottom), plotted against the physical distances (in well units) for both linear and circular mazes. Shown are the means (filled triangle and circle) ± s.e.m. (errorbars). *$p < 0.05$, **$p < 0.01$ by Wilcoxon sign-rank test. Note that these plots largely resemble the geodesic distances plots shown in Figure 5D, where only 5 and 6 trial components were used for HPC and MEC, respectively. **(J)** Average number of errors (licks at incorrect wells) made by the animals per well-combination in individual mazes for each paired session (colored diamonds and circles). Task performance was similar, in spite of the change in maze geometry. Open diamonds and circles represent the means.

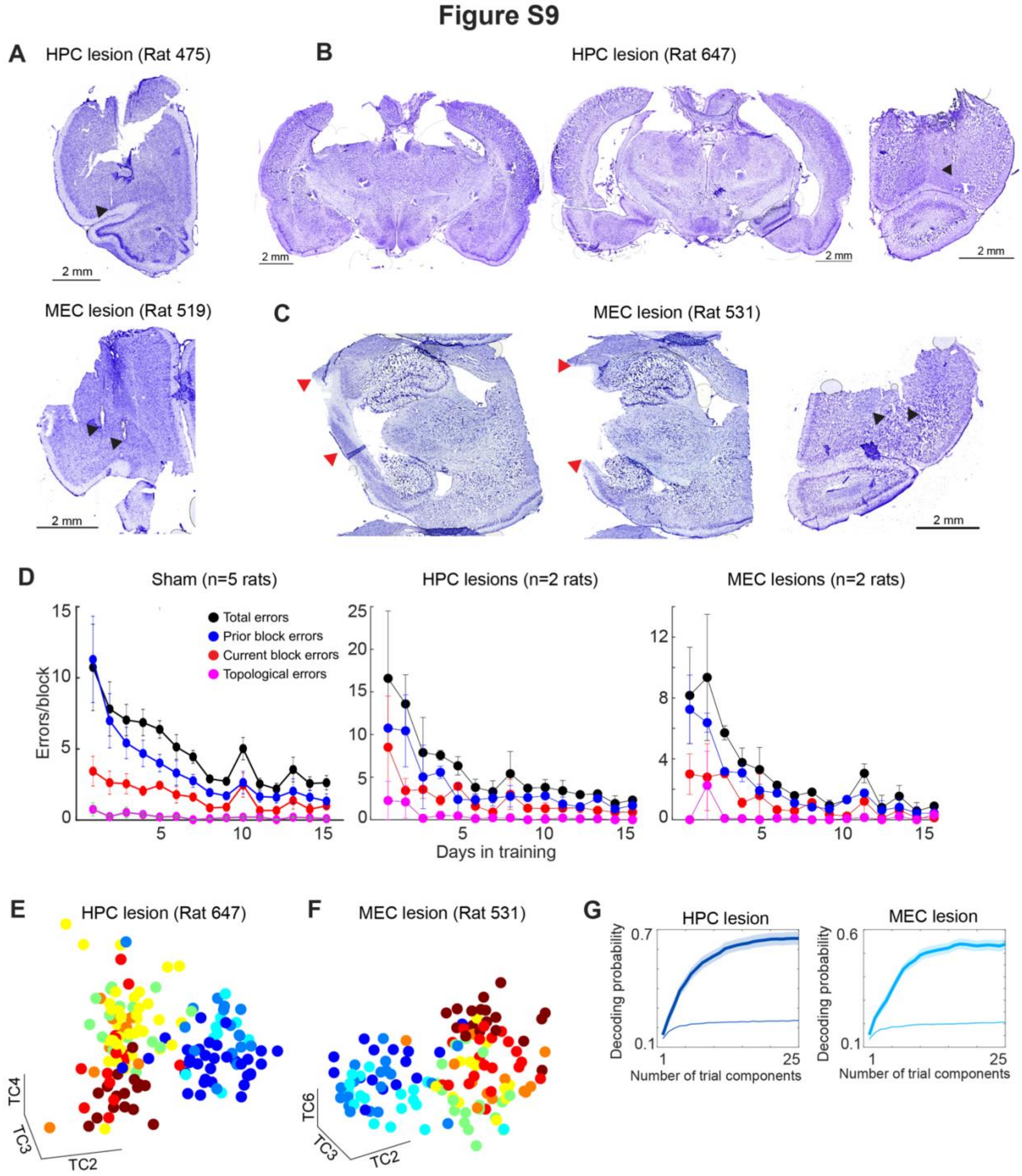


**Figure S9: OFC spatial map formation and task performance were unaffected by lesioning of HPC and MEC (related to Figure 7): (A)** Nissl-stained sections highlighting the tetrode locations targeting the OFC in the example animals shown in Figures 7A and 7B. **(B-C)** Histological sections from another set of HPC (B) and MEC (C) lesion animals. Left and middle panels show the lesions, while the right panels show the tetrode marks in the OFC. In panels S9 A-C, tetrode positions are highlighted with black arrowheads. **(D)** Learning curves of sham, HPC lesion, and MEC lesion animals. Left panel is adapted from Basu et al. 2021. Total errors correspond to the average number of errors performed per block. Prior block errors are erroneous licks of wells that were rewarded in the previous block. Current block errors constitute erroneous licks of a reward well after the reward has already been consumed at that location before visiting the paired well. Topological errors are defined as the erroneous licking of a well that is adjacent to the correct goal well (as long as the adjacent well is not rewarded in the prior block). Animals from both lesion groups follow a learning trajectory similar to that of sham animals. **(E-F)** Plots of three TCA trial components illustrating the topology preserved mapping of target wells in a representative session each from the animals with lesions in either HPC (E) or MEC (F). The examples shown are from different animals than those shown in Figures 7E and 7F. **(G)** Decoding probabilities of target well plotted as a function of the number of TCA trial components. Solid lines with shading represent means ± s.e.m. (n = 11 sessions for HPC lesions and 12 sessions for MEC lesions). Similar to Figure 2G, eight trial components can achieve 81.07% and 84.28% of the saturating decoding levels in HPC lesions and MEC lesions, respectively.
